# Neuraminidase-on-a-string nanoparticles probe how antigenic distance shapes elicited humoral immunity

**DOI:** 10.1101/2025.08.21.671615

**Authors:** Rochel Hecht, Faez Amokrane Nait Mohamed, Daniel Lingwood, Aaron G. Schmidt

## Abstract

Understanding how antigenic distance influences cross-reactive responses can inform vaccine design. Multivalent displays of viral proteins can improve B cell activation due to receptor cross-linking, and mosaic nanoparticles that incorporate variants can lead to cross-reactive B cell responses recognizing conserved epitopes. Here, we used the influenza virus neuraminidase to develop a neuraminidase-on-a-string platform displaying neuraminidase dimer pairs conjugated to a nanocarrier To systematically assess the influence of antigenic distance on humoral immunity, we paired H2N2 neuraminidase with either divergent H3N2 or H11N9 neuraminidases. We found that nanoparticle immunizations with heterologous antigens elicited sera with greater breadth and enhanced enzymatic inhibition relative to immunizations that incorporated a single neuraminidase strain. While sera reactivity for H2N2 neuraminidase was not impacted by inclusion of a second strain, strain-specific responses correlatively increased with the antigenic distance between neuraminidase components. These data show how neuraminidase strain selection for multivalent display immunizations influences elicited breadth and cross-reactivity, highlighting findings that may extend to other viral antigens.

## INTRODUCTION

Humoral responses directed at surface-exposed viral proteins (*e.g.*, coronavirus spike and influenza hemagglutinin) drive viral evolution and escape^1–3^. However, viruses cannot readily mutate conserved regions without impairing viral fitness^4^. A goal of rational immunogen design strategies, such as hyperglycosylation, epitope grafting and multivalent display aim to focus humoral immunity toward conserved epitopes to limit viral escape^5–10^. Multivalent immunogen displays increase B cell receptor crosslinking and subsequent activation^15^. Indeed, *in vivo* studies comparing monomeric versus multimeric antigens show increased antibody titers, B cell activation, and T cell engagement with multimerization^16–24^. Mechanistically, *in vivo* studies also showed that restriction to germinal center (GC) entry imposed by B cell affinity and precursor frequency thresholds are mitigated through increased avidity on an immunogen^25^. The lowered thresholds for GC entry may, in turn, increase B cell clonal diversity and breadth^22^. The size of multivalent immunogens is important, as larger particles induce slower trafficking within the lymph node^20,22,23^, leading to a prolonged antigen exposure and duration of GCs^18^. Thus, multivalent display, including viral-like particles^26^, self-assembling protein nanoparticles^27^, and DNA-origami^15^, are promising platforms for next-generation viral vaccines^20^.

Mosaic displays of distinct viral protein variants on the same nanoparticle (NP) enhanced breadth and neutralization relative to NP displays of a single protein^28–30^. This is likely due to a selective advantage for B cell receptors that engage conserved epitopes shared between the displayed antigens relative to the receptors that engage less conserved regions. However, further studies are needed to clarify the mechanisms involved, such as whether a minimum level of diversity is required to promote this advantage and whether significant variation between antigens might elicit strain-specific responses at the expense of cross-reactive responses. For example, mosaic hemagglutinin NPs showed increased proportions of cross-reactive B cells using mosaic displays only when including 6 or more strains on the same NP^29^. Additionally, whether mosaic displays stimulate cross-reactivity better than a homologous “cocktail” of the same strains has yet to be clarified. Indeed, studies incorporating antigenically equivalent cocktail versus mosaic immunogens have indicated mixed outcomes in terms of benefits^29–33^. Moreover, for both mosaic and homologous cocktail experiments, data indicate that different combinations of 2 or 4 heterologous strains resulted in different sera cross-reactivity profiles^28,34^.

Here, we addressed the potential mechanism(s) underlying these observations, and to understand how antigenic distance between displayed proteins on a NPs influenced elicited sera breadth and cross-reactivity. We chose the influenza neuraminidase (NA) as our prototypic antigen. NA catalyzes the cleavage of terminal sialic acids, and its function is essential for influenza viral egress and spread^35–37^. There is growing interest in NA as a vaccine target^38–41^, as antibodies targeting NA can be broadly reactive and protective against influenza infection^42–44^. Thus, inclusion of an NA component in next-generation influenza vaccines may provide an independent arm of immune protection against influenza, contributing to seasonal vaccine effectiveness^45,46^.

Briefly, we engineered a series of dimeric NA head constructs tandemly linked by a rigid peptide to create neuraminidase-on-a-string (NoaS) immunogens, which were then conjugated to ferritin NPs via SpyTag/SpyCatcher^47^. Each dimer contained the NA from H2N2 Japan 1957, but the second NA component was a distinct strain, which varied the antigenic distance up to ∼50% between the NA pairs. Homologous dimers of each NA were used as controls. The dimeric design enabled consistent molar ratios and spatial orientations of the NA components. Mice were immunized with either heterologous (*i.e.,* mosaic) NoaS NPs or an equivalent equimolar homologous cocktail of NoaS NPs. Serum samples were collected before and after boosting and evaluated for anti-NA breadth, strain-specificity, and NA enzymatic inhibition. We found that including a second NA strain in immunizations enhanced both anti-NA breadth and NA enzymatic inhibition. Notably, the proportion of strain-specific antibodies for each NA component increased with the antigenic distance of the NoaS pairing, even as overall anti-NA reactivity remained comparable. These findings guide next-generation viral vaccine development and underscore the importance of strain selection in multivalent vaccine platforms designed to maximize elicited cross-reactive immune responses.

## RESULTS

### Design of neuraminidase-on-a-string nanoparticles

We selected four influenza neuraminidase (NA) strains to be incorporated in our immunogens: H2N2 A/Japan/305/1957 (J’57), H3N2 A/Bilthoven/21438/1971 (B’71), H3N2 A/Darwin/6/2021 (D’21), and H11N9 A/tern/Australia/G70C/1975 (Au’75). We chose these strains primarily based on their amino acid diversity relative to the J’57 NA head: B’71, D’21, and Au’75 differed from J’57 by ∼8%, ∼17%, and ∼50%, respectively (**Fig. 1A,B** and **fig. S1C**). Additionally, the H2N2 as well as both H3N2s previously circulated within the human population, while H11N9 represents a potentially zoonotic influenza that is within the avian population. To understand how antigenic distance between NAs on a multivalent nanoparticle (NP) immunogen influenced sera cross-reactivity and breadth, we designed neuraminidase-on-a-string (NoaS) NPs. We tandemly linked pairs of monomeric NA heads together using a rigid, proline-rich L3 amino acid peptide linker^48^ to make NoaS and subsequently conjugated them to the *Helicobacter Pylori* ferritin NP^49^ using SpyTag/SpyCatcher^47^ (**Fig. 1B, C** and **fig. S1A-B**). We designed homodimeric NoaS NPs of each of these four strains (groups 1, 2, 4, and 6; **Fig. 1C**) and made heterodimeric NoaS NPs by pairing J’57 with either B’71, D’21, or Au’75 (groups 3, 5, and 7; **Fig. 1C**). Thus, using ‘percent amino acid difference’ as a proxy for antigenic distance, we varied the presented antigenic distance on the NoaS NPs from 0% up to ∼50%. Each NoaS NP displayed a total of 48 copies of the NA head, with one NoaS dimer per ferritin protomer. In the heterologous NoaS, the J’57 NA strain was conjugated adjacent to the ferritin protomer with the second NA component projecting outwardly (**Fig. 1C**).

**Fig. 1.**
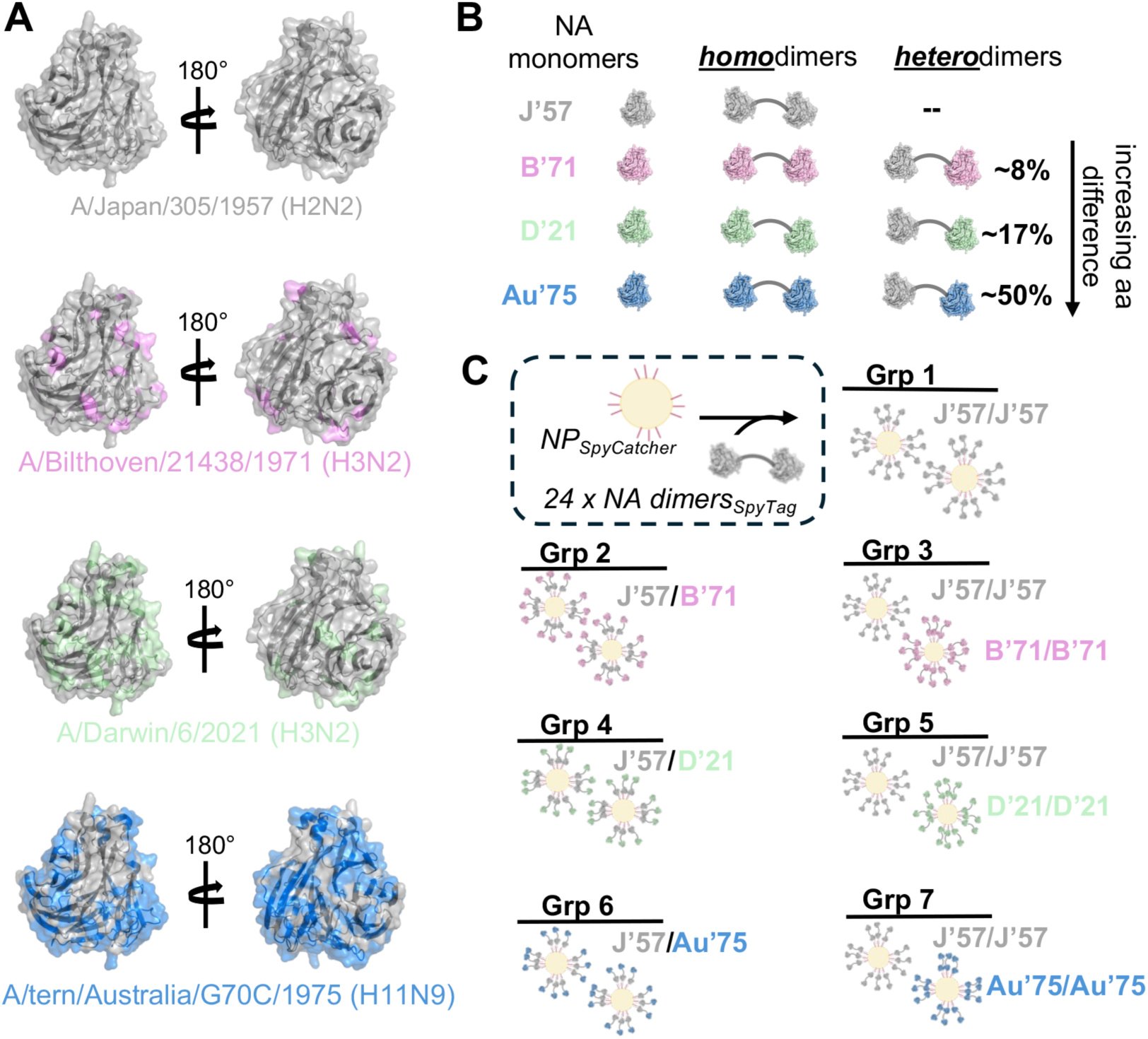
Design of neuraminidase-on-a-string nanoparticles. **(A)** The four neuraminidase (NA) strains included in the experiment were A/Japan/305/1957 (J’57), A/Bilthoven/21438/1971 (B’71), A/Darwin/6/2021 (D’21), and A/tern/Australia/G70C/1975 (Au’75). A monomer cartoon of the J’57 strain (PDB 6Q20^42^) is shown in gray, with a direct view of the catalytic site. Amino acid residues that differ between J’57 and B’71, D’21, and Au’75 NA strains are highlighted in pink, green, and blue, respectively. **(B)** Schematic of the homo- and heterodimer NA antigens. A rigid “L3” protein linker^48^ (gray curved line) separates the NAs in each dimer. Approximate percent identity amino acid difference is noted for the between the NA monomers in the heterodimers; this is relative to the J’57 NA sequence. **(C)** The NoaS NP groups used *in vivo* are shown schematically. 24 NA dimers (48 total NAs) are conjugated via a SpyTag to SpyCatcher on the N-termini of the ferritin protomers. J’57/J’57 NA homodimer (Group 1) is the control immunogen. Groups 2, 4, and 6 consisted of an equimolar cocktail of homologous J’57/J’57 NPs with B’71/B’71, D’21/D’21, or Au’75/Au’75 NPs, respectively. Groups 3, 5, and 7 display 24 copies of NoaS heterodimers on each NP. Each heterodimer contained two strains: J’57 and B’71 (Group 3), D’21 (Group 5), or Au’75 (Group 7).

### Biochemical characterization of NoaS NPs

NoaS and NoaS NPs were recombinantly expressed from mammalian cells and purified with affinity and size exclusion chromatography (SEC). The C-terminal affinity tags were removed by enzymatic cleavage and re-purified by SEC. The NoaS, in molar excess, were then conjugated to the NPs and purified again over SEC. The resulting NoaS NPs were monodisperse (**Fig. 2A and fig. S2B**) and homogeneous as assayed by SDS-PAGE analysis (**Fig. 2B**). All NoaS NPs were tested for reactivity to conformation-specific monoclonal antibodies (mAbs) NA73^50^, 3A10^51^, 1G01^42^, and NDS.1^52^ to assess structural integrity of the NA components (**Fig. 2C and fig. S2A**). NoaS dimers and NoaS NPs had comparable reactivity to the diagnostic mAbs, indicating that each NA component was in its native conformation and accessible after conjugation to the ferritin NP. NoaS NPs were visualized using negative stain electron microscopy and showed uniform NoaS NP sizes of ∼60nm, corresponding well to the approximate additive size of all components (**Fig. 2D**). The L3 linker^48^ spaced each of the ∼5nm-wide NA heads about 10nm apart (**fig. S2C**). This approximate distance between the two NA heads is optimal to engage both antibody arms of a B cell receptor (∼13nm distance)^53^.

**Fig. 2.**
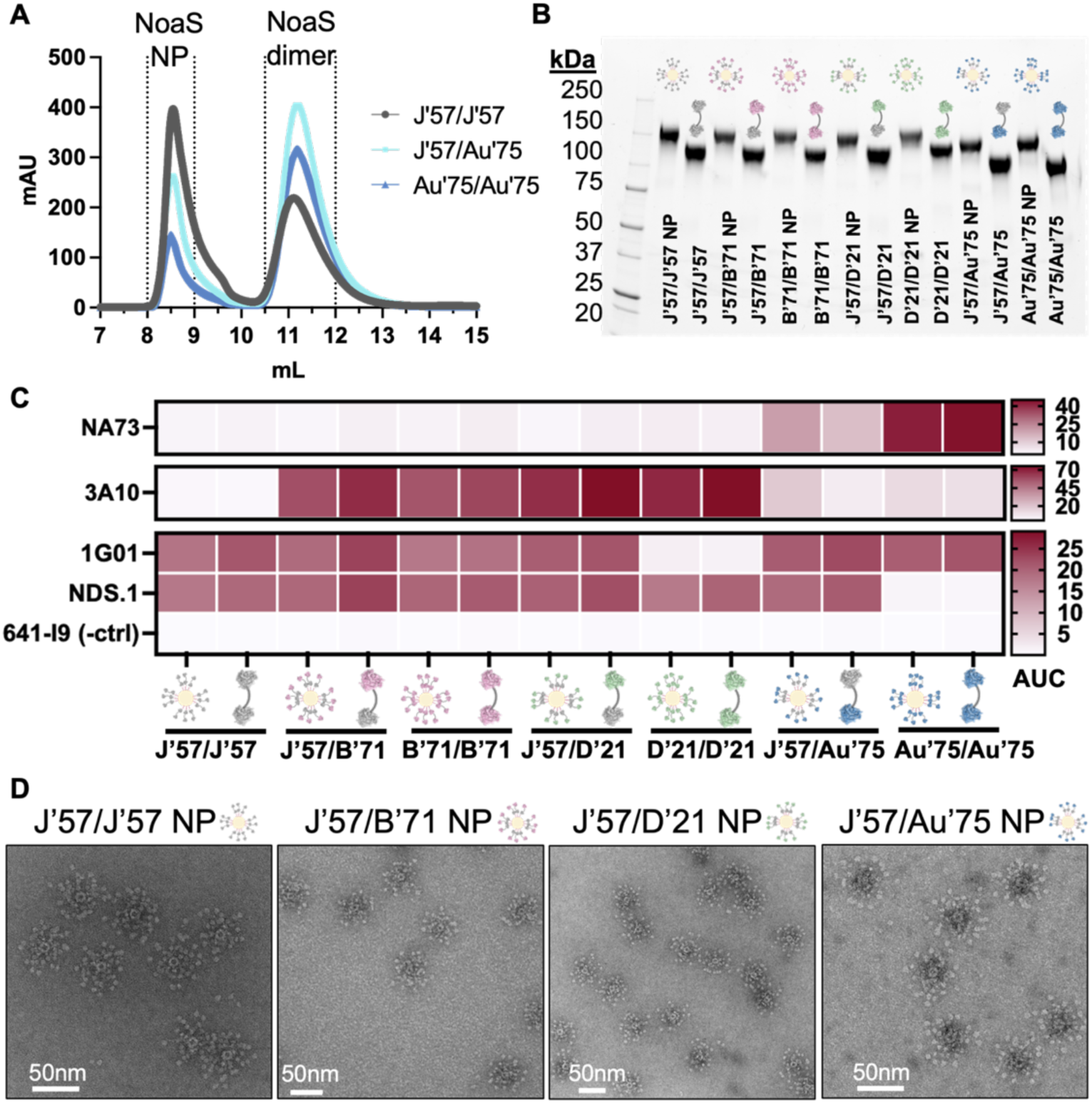
Biochemical characterization of NoaS and NoaS NPs. **(A)** Size exclusion chromatography traces of NoaS NPs with excess NoaS dimers after conjugation. **(B)** SDS PAGE with NoaS dimers conjugated to NP protomers and NoaS dimers. (Note: ferritin NP is non-covalently assembled and dissociates into protomers in SDS-PAGE). **(C)** Reactivity of strain- and conformation-specific mAbs to NoaS NP. NA73 mAb is N9-specific, used for Au’75 NA detection. NDS.1 mAb binds J’57, B’71, and D’21 NAs but does not bind Au’75 NA. It was used in combination with NA73 mAb to verify the J’57/Au’75 NA components. 3A10 mAb binds B’71 and D’21 NAs but does not bind J’57 NA. 1G01 mAb does not bind D’21 NA. 3A10 mAb was used with 1G01 mAb to detect J’57/D’21 NAs. 3A10, NDS.1, and 1G01 mAbs detected J’57/B’71 NAs. 641-I9 mAb^54^ is influenza HA-specific and was used as a negative control. **(D)** Representative negative stain micrographs of NoaS NP from each group.

### Group design and vaccine regimen

Seven groups of C57BL/6 mice (n=5 per group) were immunized using a homologous prime-boost regimen (**Fig. 3**). Mice received 100μL of inoculum intraperitoneally, containing 20μg of total protein adjuvanted with 50% w/v Sigma adjuvant. Group 1, which received J’57/J’57 NoaS NPs, served as the control group. Groups 3, 5, and 7 received heterologous NoaS NPs displaying J’57 with either B’71, D’21, or Au’75, respectively. Groups 2, 4, and 6 received a cocktail of homologous J’57/J’57 NPs with either B’71/B’71, D’21/D’21, or Au’75/Au’75 NPs, respectively. The cocktails were prepared in equimolar ratios to their heterologous counterparts. Mice were bled weekly, and the sera were collected 14-27 days post-prime and 41-62 days post-prime (*i.e.,* 14-35 days post-boost) to be evaluated for NA breadth, NA strain-specificity, and NA enzymatic inhibition.

**Fig. 3.**
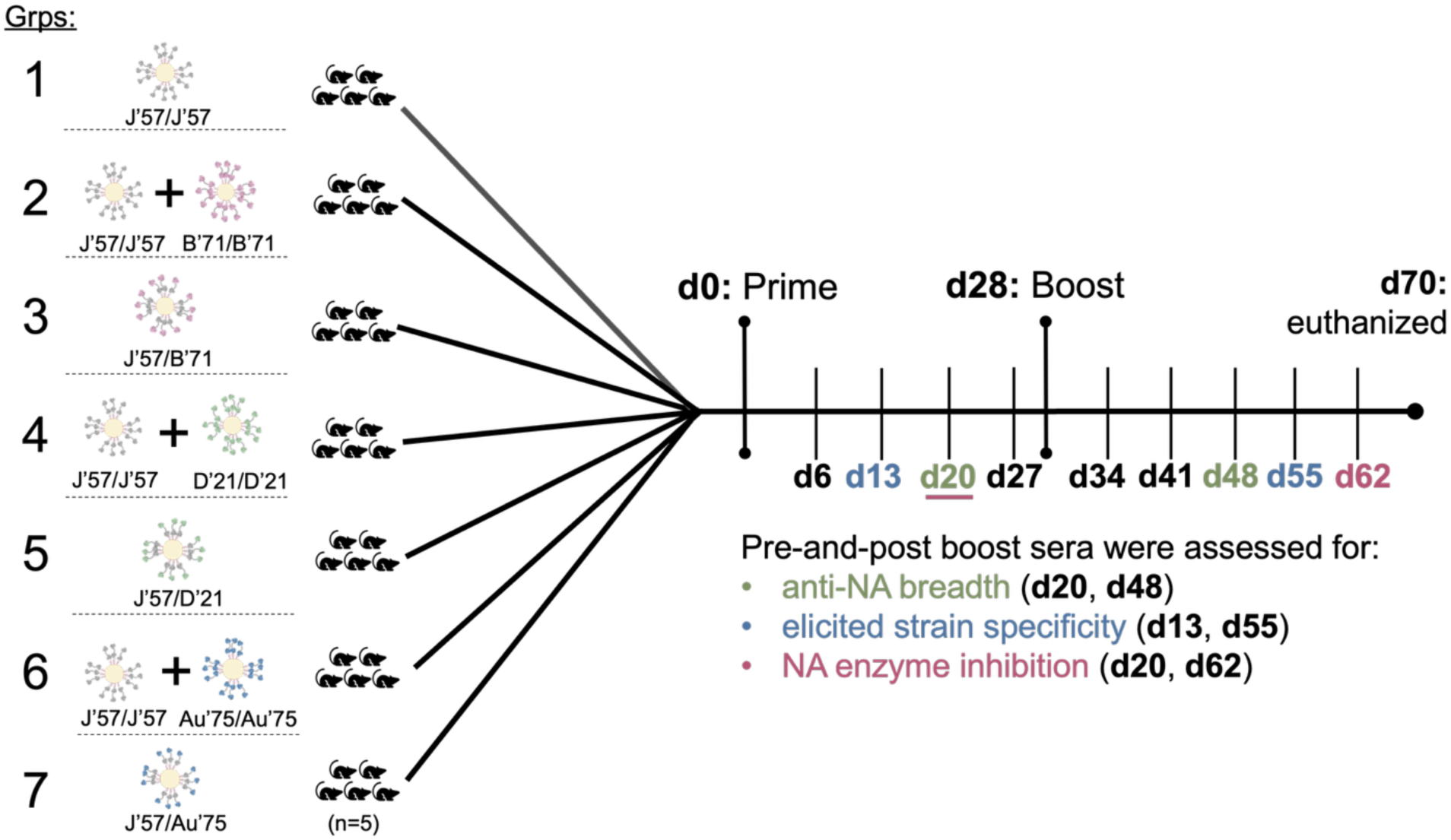
NoaS NP groups and immunization and assessment schema. Seven groups of C57BL/6 mice (n=5 per group) were immunized intraperitoneally on day (d) 0 and d28 and euthanized on d70. Serum from each mouse was collected weekly. Sera were evaluated on either the second or third week after the prime and up to 5 weeks after the boost for: breadth elicited against NA strains, degree of developed strain-specificity, and ability to inhibit NA enzymatic activity.

### Assessment of elicited sera breadth in NoaS NP groups

A panel of six historical N2 heads (A/Japan/305/1957 (J’57), A/Bilthoven/21438/1971 (B’71), A/Aichi/2/1968 (Ai’68), A/Moscow/10/1999 (M’99), A/Texas/50/2012 (Tx’12), and A/Darwin/6/2021 (D’21)) and one H11N9 (A/tern/Australia/G70C/1975 (Au’75)), which included the four NA strains incorporated into the immunogens, were recombinantly expressed as monomers. The amino acid differences between the NA strains in the panel varied from ∼4.4-53.5% (**fig. S3A**) and the number of predicted-N-linked glycosylation sites in the panel ranged from 3-7 (**fig. S3B**). Sera from d20 and d48 were tested for reactivity to the entire NA panel in ELISA. Absorbance values were normalized per antigen, and the degree of normalized area under the curve (nAUC) was plotted for each mouse serum sample (**figs. S4A-B** and **S5A-B**). Except for the reactivity of group 4 versus group 5 to the Tx’12 antigen on d20 (**fig. S6B**), there were no statistically significant differences in the observed reactivity to the NA panel when comparing the homologous cocktail groups (*i.e.,* groups 2, 4, and 6) to their heterologous NoaS counterparts (*i.e.,* groups 3, 5, and 7) (**fig. S6A-C**). This suggested that any effect of heterologous strain presentation versus homologous cocktail presentation was likely too small to be observed with our n=5 group size. We therefore combined the data from groups 2 and 3, groups 4 and 5, and groups 6 and 7, respectively, to probe how the antigenic distance within a two-strain immunization influenced elicited breadth. (Note: In this study, *serum breadth* refers to the overall extent of unique NA strains recognized by the serum in our NA panel. *Serum cross-reactivity* describes when individual antibodies within the serum recognize more than one antigenically distinct NA strain. Serum breadth can result either from cross-reactive serum antibodies (*i.e.,* the same serum antibodies binding multiple NA strains) or from a more diverse serum antibody pool (*i.e.,* different serum antibodies recognizing different NA strains). Thus, while serum that recognizes distinct NA strains has greater breadth than serum recognizing only one strain, this does not necessarily imply that it contains more cross-reactive serum antibodies.)

We found that all groups had comparable reactivity to the J’57 antigen (**Fig. 4A**), which indicated that J’57 immunogenicity was not affected by the inclusion of a second NA strain. Additionally, all experimental groups had greater anti-NA breadth than the control group, implying that inclusion of a second strain increased the elicitation of NA breadth (**Fig. 4B-G**). On d20, J’57/D’21 groups had the broadest reactivity across the tested N2 NA panel (**Fig. 4C-F**). On d48, although the groups receiving J’57/D’21 still showed broad N2 NA reactivity, the groups receiving J’57/B’71 developed statistically significant reactivity to earlier N2 NA strains (*i.e.,* Ai’68 and B’71; **Fig. 4B-C**). These data suggested that the inclusion of two antigenically similar NA strains (*i.e.,* J’57 and B’71) could enhance the reactivity to each other as well as to strains antigenically like each other (*i.e.,* Ai’68). Finally, the groups receiving J’57/Au’75 had N2 NA breadth that was comparable to the control group across the N2 panel but also had reactivity to the N9 strain included in the NoaS (**Fig. 4A-G)**. These data suggested that the antigenic distance between J’57 and Au’75 was too great to elicit broader N2 reactivity. Nonetheless, including the N9 NA allowed strain-specific recognition of each component.

**Fig. 4.**
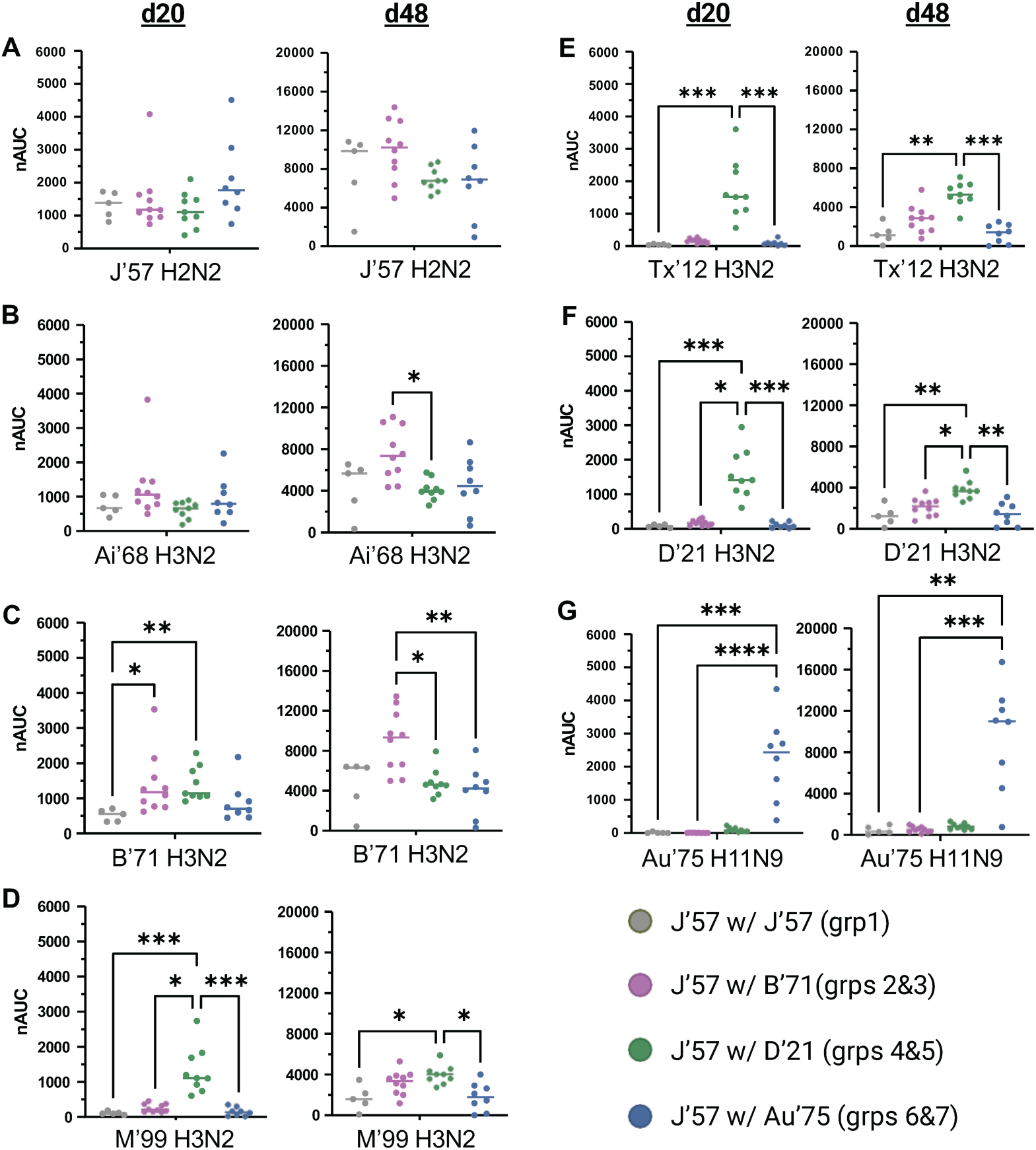
NA breadth of d20 and d48 sera. **(A)** Sera reactivity to the J’57 NA monomer. No statistically significant differences in reactivity were observed. **(B)** Sera reactivity to the Ai’68 NA monomer. Statistically greater reactivity to the Ai’68 NA strain on d48 is noted for groups receiving J’57/B’71 NoaS NPs over groups receiving J’57/D’21 NoaS NPs (*0.0122). **(C)** Sera reactivity to the B’71 NA monomer. On d20, groups that received either B’71 or D’21 NA as their second NA component in immunizations had statistically greater reactivity to B’71 NA over the control group (*0.0224, **0.0064). On d48, groups that received B’71 NA had greater reactivity to the B’71 NA than the remaining experimental groups (*0.0150, **0.0052). **(D)** Sera reactivity to the M’99 NA monomer. Groups receiving D’21 NA in their immunizations had statistically greater reactivity to the M’99 NA on d20 (*0.0371, ***0.0005 for control, ***0.0003 for Au’75) and on d48 (*0.0203 for control, *0.0110 for Au’75). **(E)** Sera reactivity to the Tx’12 monomer. On both d20 and d48, groups that received D’21 NA in immunizations had statistically greater reactivity to the Tx’12 NA strain than the control group or groups receiving Au’75 NA (d20: ***0.0001 for control, ***0.0004 for Au’75. d48: **0.0023, ***0.0004). **(F)** Sera reactivity to the D’21 NA monomer. Groups receiving D’21 NA in their immunizations showed statistically greater reactivity to D’21 NA than all other groups on both d20 and d48. (d20: ***0.0007 for control, *0.0279, ***0.0004 for Au’75. d48: **0.0037 for control, *0.0368, **0.0018 for Au’75.) **(G)** Sera reactivity to the Au’75 NA monomer. Groups that received Au’75 NA in their immunizations had statistically greater reactivity to the Au’75 NA strain than other groups. (d20: ***0.0003, ****<0.0001. d48: **0.0043, ***0.0009.) Statistical differences were assessed using unpaired, non-parametric Kruskal-Wallis tests with Dunn correction for multiple comparisons. Horizontal lines represent the median. Data from mice (m) m21, m28, and m31 were removed from all statistical analyses due to lack of response to the prime immunization.

To assess whether serum antibodies from the groups acquired breadth by focusing on the conserved NA catalytic site (CS), we tested sera reactivity to a modified J’57 NA with a glycan projecting into the CS. This ‘ΔCS’ J’57 version abrogated binding to all CS-directed mAbs assayed (**fig. S7A**). By comparing the sera reactivity to the wildtype J’57 versus ΔCS J’57 antigens, we could compute the percent loss of signal due to the glycan and assess how much of the sera was directed at or near the CS. We did not observe any differences in ΔCS J’57 reactivity in the cocktail versus heterologous groups, so we analyzed groups based on the pair of NA strains received in the immunizations (**fig. S7B-C**). On both d20 and d48, there were statistically comparable reactivities to the ΔCS J’57 antigen across all paired groups (**fig. S7D**). On d20, reduced reactivity to the ΔCS J’57 antigen compared to the wildtype J’57 antigen was significant for the J’57/J’57, J’57/B’71, and J’57/Au’75 groups. By d48, the reduced reactivity was significant for all groups except J’57/J’57, however this group trended similarly (**fig. S7E**). These results suggest that, across groups, the inclusion of a second, antigenically distinct NA strain contributed to focusing the sera responses on the CS, which is conserved across NA strains.

Lastly, scaffold-specific sera responses to the ferritin NP were assayed on d27 and d62 (**fig. S8A-C**). On d27, the NP reactivity was similar across groups. However, on d62, the J’57/B’71 and J’57/Au’75 had enhanced reactivity toward the naked NP relative to the J’57/J’57 control group (**fig. S8C**).

### Strain-specificity of elicited sera responses

We developed a serum depletion assay to evaluate whether NA pairs elicited strain-specific or cross-reactive antibodies (**fig. S9-19**). Each serum sample was first tested by ELISA for reactivity against both NA components used in each immunization. For example, serum from mouse 30 (m30) in group 6 (which received both J’57 and Au’75 NAs) was first assayed for binding to J’57 and Au’75 NAs (**fig. S12A**). The m30 serum was then incubated over a column containing J’57 NA, thereby depleting the sample of J’57- reactive antibodies. The J’57-depleted serum was subsequently tested for reactivity to both J’57 NA (**fig. S12B**) and Au’75 NA (**fig. S12C**) by ELISA. The absence of any remaining J’57 NA reactivity confirmed successful depletion, and the remaining Au’75 NA reactivity was compared to the pre-depletion level. The ratio of Au’75 NA reactivity post- vs. pre-J’57 depletion reflected the degree of strain-specificity for Au’75 (**fig. S13E**).

Importantly, this assay quantified how much of the serum antibody response to a given NA was specific to that NA. It did not directly measure the percentage of antibodies that were cross-reactive. For instance, a strain-specificity value of ∼79% for the Au’75 NA in m30 did not imply that ∼21% of total serum antibodies were cross-reactive. Rather, it meant that *of* the antibodies binding Au’75 NA, ∼21% also recognized J’57 NA. In this way, the assay measured how much of the response to strain A was unique to strain A, and it only indirectly reflected cross-reactivity with strain B. Using this approach and analysis, sera-depletion assays were performed on each serum sample collected on d13 and d55.

The homologous cocktail groups 2, 4, and 6 had similar strain-specificity profiles for their counterpart heterologous groups 3, 5, and 7 on both d13 and d55 (**fig. S20A-B**). We therefore analyzed data from groups 2 and 3, 4 and 5, and 6 and 7 as combined cohorts, respectively. The J’57/J’57 group acquired some breadth for B’71, D’21, and Au’75 NAs (**fig. S22A-B**). However, after sera depletion for J’57-reactive antibodies, the remaining sera no longer recognized B’71, D’21, and Au’75 NAs (**figs. S13** and **S18**). Thus, we conclude that none of the serum antibodies elicited in the J’57/J’57 group were strain-specific for B’71, D’21, or Au’75 NAs.

The remaining groups were depleted for both J’57 NA reactivity and the second NA component in their immunogens (**figs. S9-S18**). On d13, all groups had comparable reactivity to J’57 NA (**Fig. 5A**), and most groups had comparable reactivity to J’57 NA on d55. Nonetheless, the degree of strain-specificity for J’57 NA increased in the groups in correlation with the antigenic distance of their second NA component (**Fig. 5B**). This same phenomenon was observed for the NA strain that J’57 was paired with (**Fig. 5C**). Thus, groups that received J’57/B’71 had substantially less strain-specificity toward the J’57 NA than did the groups that received J’57/Au’75. Similarly, the calculated strain-specificities toward B’71 NA in groups 2 and 3 were substantially lower than the calculated strain-specificities toward Au’75 NA in groups 6 and 7. Thus, pairing the J’57 NA with an NA strain that had increased antigenic distance elicited a greater proportion of serum antibodies specific to each NA component. When comparing the strain-specificity on d55 versus d13, all groups showed a decrease in the magnitude of specificity for both the J’57 NA (**fig. S21A**) and their paired NA component (**fig. S21B**), with varying statistical significance across groups.

**Fig. 5.**
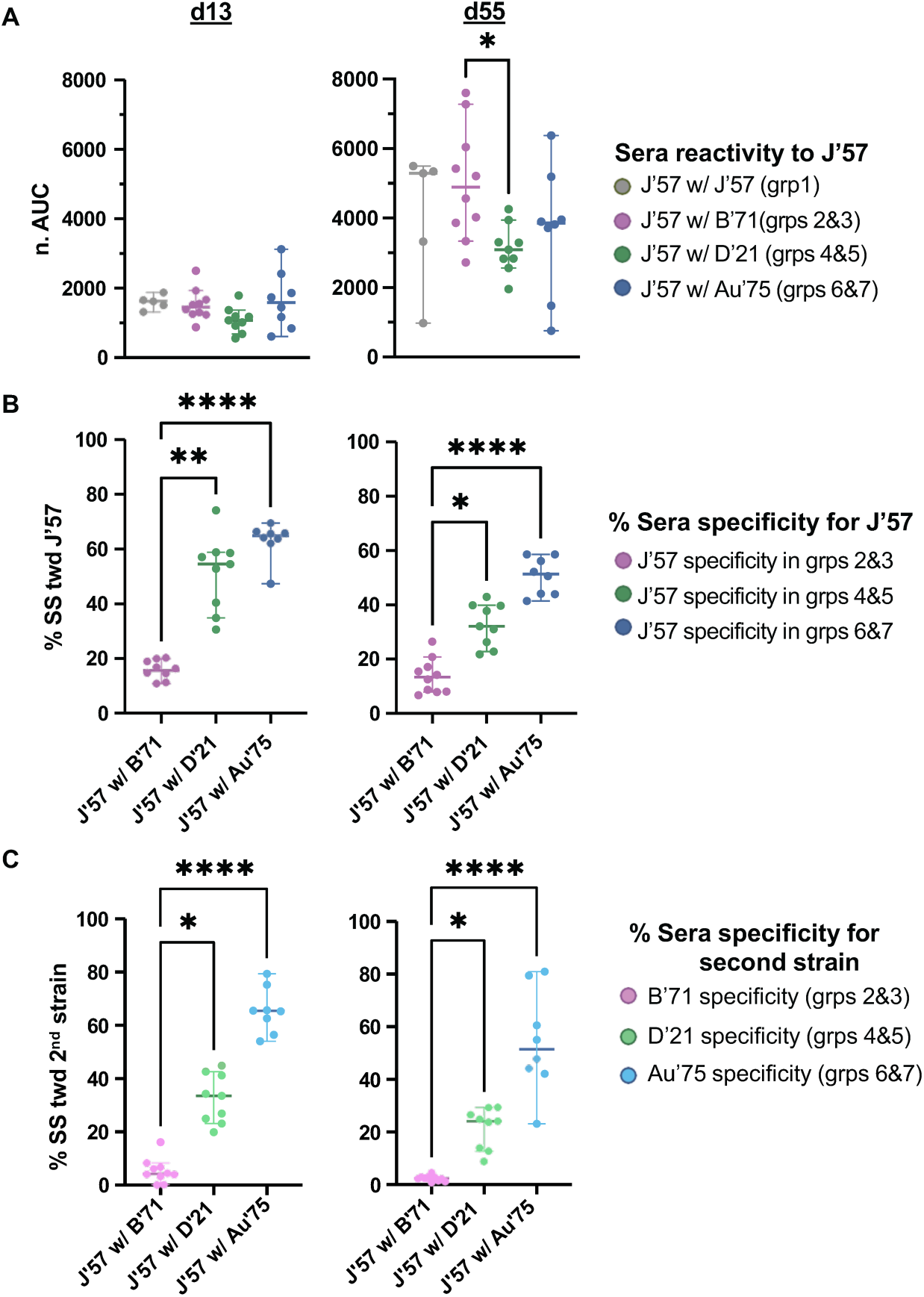
Strain-specificity toward J’57 NA and the second NA component. **(A)** d13 and d55 sera reactivities toward the J’57 NA. On d13, reactivities toward the J’57 strain were not statistically different across groups. On d55, reactivity toward the J’57 strain was significant between groups receiving B’71 NA versus D’21 NA as their second NA component (*0.0408). **(B)** Strain-specificity for the J’57 NA on d13 (p-values: **0.0081 and ****<0.0001) and d55 (p-values: *0.0357 and ****0.0001). **(C)** Strain-specificity for the second NA component on d13 (p-values: *0.0276 and ****<0.0001) and d55 (p-values: *0.0213 and ****<0.0001). Differences in groups were assessed using the Kruskal-Wallis test with Dunn correction for multiple comparisons. Lines and bars represent the median with a 95% confidence interval.

### Sera inhibition of NA enzymatic activity

We next assessed whether the elicited sera from the NoaS NPs inhibited NA enzymatic activity using the enzyme-linked lectin assay (ELLA), which uses a large fetuin substrate^55^. Antibodies that bind directly to the CS block enzymatic activity even with small substrates, but antibodies that bind proximal to the CS or on the NA periphery can sterically obstruct the CS. The ELLA inhibition assay is therefore considered more reflective of the broader potential to inhibit NA activity over other assays that use smaller substrates, ^42,52^, such as NA-Star^TM^ or MUNANA^56^. We assayed d20 and d62 sera for enzymatic inhibition of the NA strains used in each immunization (**figs. S23-S24**). We first determined the EC_50_ concentration for a given NA strain and then incubated two times the EC_50_ with two times the serial serum dilution and measured the resulting NA activity. Using d0 serum as a negative control, we identified the dilution factor at which a given serum sample inhibited NA enzymatic activity more than the highest concentration of d0 serum (*i.e.,* the maximum serum dilution factor with inhibitory activity, or MDFI). Homologous cocktail groups 2, 4, and 6 showed similar inhibition profiles to their counterpart heterologous groups 3, 5, and 7, respectively (**fig. S25A-B**). Groups were therefore evaluated based on immunizing strains.

All groups had comparable enzymatic inhibition of J’57 NA activity on both d20 and d62, indicating that the inclusion of a second NA strain did not reduce serum antibodies with J’57 CS-inhibiting activity (**Fig. 6A** and **figs. S23A** and **S24A**). We then tested B’71 NA enzymatic inhibition in groups 2 and 3, D’21 NA enzymatic inhibition in groups 4 and 5, and Au’75 NA enzymatic inhibition in groups 6 and 7 (**figs. S23B- D** and **S24B-D**). Each NA strain was also tested against sera from the control, group 1. Sera from d62 showed substantially stronger inhibition of B’71 in groups 2 and 3 compared to the control group (**Fig. 6B**). Surprisingly, at both time points, despite the greater antigenic distance between D’21 and J’57 NAs than the B’71 and J’57 NAs, inhibition of D’21 NA activity was not significantly improved in groups 4 and 5 sera compared to the control group sera (**Fig. 6B**). Lastly, sera from groups 6 and 7 inhibited the Au’75 NA activity substantially better than sera from the control group (**Fig. 6B**). These data show that including a second NA component in NoaS NPs with J’57 NA does not detract from J’57 NA activity inhibition and, moreover, better ensures the production of antibodies with inhibitory potential to a second NA.

**Fig. 6.**
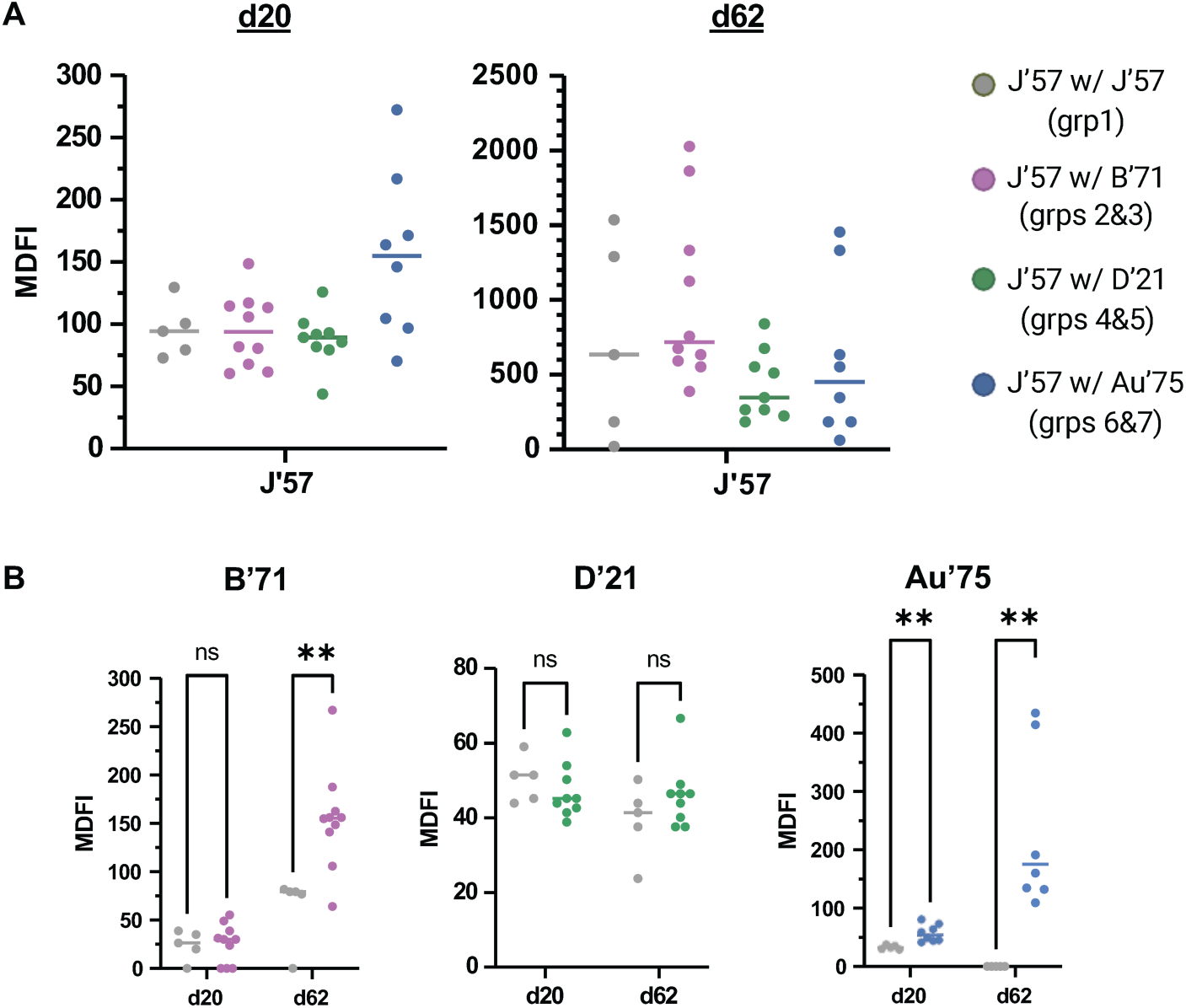
Maximum dilution factor of inhibition (MDFI) of sera as assessed in ELLA inhibition assays. **(A)** ELLA inhibition of J’57 NA. Sera from all groups showed statistically comparable inhibition of J’57 NA activity on d20 and d62. Data were analyzed using the Kruskal-Wallis test with Dunn corrections for multiple comparisons. **(B)** ELLA inhibition on d20 and d62 of B’71, D’21, and Au’75 NAs. Statistical significances are noted (B’71 groups: **0.007, Au’75 groups: **0.002). Data were assessed using Mann-Whitney tests. Lines reflect the median MDFI values. Serum from d0 was used as the negative control.

## DISCUSSION

Here, we engineered a multivalent, modular neuraminidase-on-a-string (NoaS) nanoparticle (NP) platform to assess the effects of varying neuraminidase (NA) antigenic distance on the elicited humoral response. Sera reactivity to the NA panel yielded two key observations. First, J’57 NA immunogenicity was not reduced when a second NA strain was included in the NoaS NPs, regardless of the antigenic distance relative to J’57 NA. Second, anti-NA breadth is augmented when a second, antigenically distinct NA strain is included. Additionally, the proportion of strain-specific serum antibodies increases with the antigenic distance between the NA components, and that this relationship remains consistent after boosting. We also found that inhibition of J’57 NA enzymatic activity by sera was not impacted by including a second NA strain in NoaS NPs; this observation may correlate with protection.

Interrogating scaffold-specific responses is necessary to ensure that next-generation vaccine platforms do not abrogate desired immune responses to the viral immunogens^57^. Since each NoaS dimers were approximately similar in size across experimental groups, we anticipated that the NoaS dimers would sterically occlude access to the ferritin surface equivalently and thus have comparable scaffold-specific responses. While this was generally observed, at d62 (*i.e.,* 35 days post-boost) the J’57/B’71 and J’57/Au’75 experimental groups elicited stronger ferritin-specific responses relative to the J’57/J’57 control group (**fig. S8C**). It is unclear why these NA pairings resulted in a more robust scaffold response however, these experimental groups also had stronger reactivity to both NA components on d20 and d48 (**Fig. 4A, C, and G**). Importantly, the observed scaffold-specific responses did not appear to reduce sera responses to the NA components. Thus, using the ferritin NP to increase multivalency is a possible platform for NA-based vaccines.

Incorporating diverse strains of the same viral protein into a vaccine can induce humoral responses to each individual strain^12,13^. This strategy of enhancing breadth through multi-strain immunization is supported by numerous studies^28–31^. Therefore, our finding that NoaS NPs elicited serum responses to each included NA component is consistent with prior observations. Notably, the combination of J’57 and D’21 NAs elicited broad reactivity to a panel of historical N2 NAs spanning >60 years of antigenic distance. The observed reactivity may be due to cross-reactive serum antibodies recognizing both J’57 and D’21 NAs, or from distinct serum antibody populations targeting each NA strain. Data from our serum-depletion assay supported the latter: we observed that serum strain-specificity increased with antigenic distance between paired NAs on the NoaS NPs, suggesting that the J’57/D’21 group achieved breadth primarily through separately affinity-matured serum antibody pools. While some cross-reactive serum antibodies may have been elicited, they appeared to represent a minority of the response, particularly in the J’57/D’21 and J’57/Au’75 groups.

In general, we did not find that the mosaic displays of the heterologous strains resulted in statistically significant greater NA breadth, cross-reactivity, or activity inhibition than their homologous cocktail counterparts. Previous studies also showed “cocktails” of individual components were generated comparable^32^ or even greater^30^ cross-reactivity within the sera than the mosaic displays. Considering our data and these discrepant studies, we suspect that it is possible (if not likely) that the benefits of enriching for conserved epitopes through the display of heterologous strains on the same NP is best stimulated using a greater number of strains while still covering a well-constrained antigenic distance^29^. While the heterologous NP groups in our study had comparable or qualitatively greater sera cross-reactivity than did the homologous cocktail groups, statistical significance might require a greater number of incorporated NA strains (*e.g.,* >2). We also observed that all groups had decreased strain-specificity at d55 (*i.e.,* 28 days post-boost) compared to d13 (**fig. S21A-B**). This observation may be due to prolonged duration of somatic hypermutation^58,59^, where antigen-reactive B cells in germinal centers generated increased cross-reactivity over time through modulating affinity for the strains incorporated in the immunization^60–62^.

Previous studies showed that NA immunizations protected against viral challenge better when the NA strain was matched rather than heterologous^63–65^. Our NA enzymatic inhibition data used as a proxy for protection are consistent with these observations: experimental groups that received B’71 or Au’75 NA strains outperformed the control J’57/J’57 group when testing for NA enzymatic inhibition against B’71 or Au’75 NAs, respectively. However, the results from groups receiving the D’21 NA, which did not inhibit D’21 NA enzymatic activity better than the control J’57/J’57 group, highlights a greater nuance. Structural analysis of D’21 and J’57 NAs suggest that it may be due to the unique glycosylation patterns on D’21 NA. D’21 NA has three different and two additional predicted N-linked glycosylation sites (PNGs) compared to the J’57 NA. In particular, the N245 and N367 PNGs are adjacent to the CS and may have therefore affected the CS-directed antibody responses to D’21 NA. Indeed, some previously characterized broadly reactive CS-directed antibodies have reduced binding to NA strains with the N245 glycan^66^. Our observations suggest that eliciting sera antibodies against specific epitopes across diverse NA strains might require additional design strategies for epitope-focusing. Rational design strategies like hyperglycosylation and epitope grafting are compatible with these multivalent NoaS NP designs and may be used r to improve humoral focusing to desired a desired NA epitope(s)^8,10^.

The immune response to homologous boosts is often dominated by antibodies generated during the prime immunization^67^, and early influenza exposures can imprint long-lasting immune biases^68^. Therefore, we specifically evaluated the NoaS NPs in an immunologically naïve cohort to understand how to elicit breadth in the context of primary imprinting. Understanding how complex immune histories, whether initiated through vaccination or infection, influence subsequent responses to the NoaS NPs is necessary. Additionally, understanding how NA antigenic distance influences single B cell responses or memory B cell formation may guide vaccine design. Nevertheless, these data show how rationally designed NoaS NP can advance our understanding of how antigenic distance influences elicited humoral immunity and lays a foundation for next-generation influenza vaccines that incorporate NA strains for improved protection against seasonal and potentially pre-pandemic influenzas.

## MATERIALS AND METHODS

### Strain Selections

The four NA strains used in the Neuraminidase-on-a-String (NoaS) nanoparticles (NPs) were A/Japan/305/1957, A/Bilthoven/21438/1971 (H3N2), A/Darwin/6/2021 (H3N2), and A/tern/Australia/G70C/1975 (H11N9). Strains were selected by performing amino acid alignments of many NA heads and finding strains that together represented a rational distribution of antigenic distances. Strains were then tested for expression as both monomers and paired NoaS dimers. In total, 20 different strains of NA and many neuraminidase-on-a-string combinations were tested. Final selections for the NoaS were based on ease of expression, distribution of antigenic distance, and structural integrity.

### Cloning of NoaS NPs

The NoaS plasmids were engineered using type IIS cloning^69^ (with a SAPI enzyme) and cloned into pVRc8400 protein expression vectors. The plasmids coded for the following: t-PA signal sequence, Histidine tag (for affinity purification), HRV 3C protease site (to cleave off the HIS tag), SpyTag sequence^47^, first NA head, L3 rigid amino acid linker^48^, and, finally, the second NA head (Fig. B.1B). The *Helicobacter Pylori* ferritin nanoparticle^49^ plasmid coded for an N-terminus Histidine affinity tag, SpyCatcher domain (**fig. S1A**), and ferritin protomer (**fig. S1A**). All nucleotide sequences were codon optimized and synthesized by IDT. The monomeric versions of these four strains of NA as well as the NA heads of A/Aichi/2/1968 (H3N2), A/Moscow/10/1999 (H3N2), and A/Texas/50/2012 (H3N2) used in the ELISA breadth assay panel, were also codon-optimized and cloned into pVRc vectors.

### Expression and purification of proteins

SpyTag NoaS dimers and SpyCatcher NP plasmids were transiently expressed with Expi293F cells (ThermoFisher). After 5-7 days, the cell culture supernatants were harvested by centrifugation. Proteins were purified by affinity chromatography using Cobalt TALON resin (TAKARA) followed by size exclusion chromatography on a Superdex 200 Increase 10/300 GL column (GE Healthcare). The NoaS Dimers were then cleaved using HRV 3C protease (ThermoFisher Pierce) with an overnight incubation at 4°C. They were eluted over Cobalt TALON resin to remove Histidine tags and HRV 3C protease, repurified over a Superdex 200, and then conjugated in 40x molar excess to previously purified SpyCatcher NPs overnight at 4°C. The conjugated NoaS NPs were purified a final time using size exclusion chromatography. (IgGs referenced in **Fig. 2C** (including the heavy- and light-chain variable domains were synthesized and codon optimized by IDT, then subcloned into pVRc8400 protein expression vectors with human constant regions. They were expressed and purified using the same workflow as the NA antigens.)

### Sodium dodecyl sulfate–polyacrylamide gel electrophoresis (SDS-PAGE)

3µg of a protein sample, diluted to a final volume of 10µL, were mixed with 10µL of non-reducing 2x Laemmli Sample Buffer (Bio-Rad, Cat#: 1610737), then boiled for 10min at 100°C. 12.5µL of sample were loaded into wells of a Mini-PROTEAN TGX Stain-Free precast gels (Bio-Rad, Cat#: 456-8026), then electrophoresed for 18-20min at 280V. Gels were imaged using a ChemiDoc (Bio-Rad) with appropriate standard imaging protocols. For the sera-depletion serum samples in **fig. S19**, 10µL of concentrated sample were used without prior standardization for protein amount.

### Negative Stain Electron Microscopy

5µl of the sample (at 5-10µg/ml) were adsorbed for 60 seconds to a carbon-coated grid (EMS www.emsdiasum.com order# CF400-CU). The grid was rendered hydrophilic via a 20 second exposure to a glow discharge (25mA). Excess liquid was removed with a filter paper (Whatman #1). The grid was then exposed briefly to a water droplet to wash away ions and salts and blotted once again on a filter paper. The grid was then stained with 0.75% Uranyl Formate (EMS catalog # 22451) for 20 seconds. After removing the excess stain with filter paper, the grids were examined in a TecnaiG² Spirit BioTWIN and images were recorded with an AMT NanoSprint43 CCD camera.

### *In vivo* immunizations

All animal experiments were approved by the Institutional Animal Care and Use Committees (IACUC) of Harvard University and Massachusetts General Hospital (Protocol #2014N000252) and were conducted in compliance with the Association for Assessment and Accreditation of Laboratory Animal Care International (AAALAC) guidelines. Mice were used at 8–10 weeks of age. The immunizations were performed using female C57BL/6 mice purchased from The Jackson Laboratory (Bar Harbor, ME). Mice were divided randomly in 7 groups of n=5 and received a prime immunization intraperitoneally on day 0, which consisted of 100uL of inoculum containing 20ug of protein (J’57/J’57 NoaS NPs as control, NoaS NPs displaying J’57 with either B’71, D’21, or Au’75 and a cocktail of homologous J’57/J’57 NPs with either B’71/B’71, D’21/D’21, or Au’75/Au’75 NPs, respectively) adjuvanted with 50% v/v Sigma adjuvant (Sigma, Cat#S6322) in sterile PBS. The boost occurred on day 28. Mice were bled from the submandibular vein for sera analyses on day -1 and then every 7 days thereafter (until day 70, when they were euthanized).

### ELISA Assays

Sera and monoclonal antibody reactivity to NA antigens were assayed by ELISA. 96-well high binding plates (Corning) were coated with 3µg/ml of NA antigens in PBS buffer at 100µl/well and incubated overnight at 4°C. Plates were blocked with 1% BSA in PBS containing 0.1% Tween-20 (PBS-T) for 1-2 hours at room temperature (RT). The blocking solution was then discarded. Plates were coated with sera or monoclonal antibody solutions and incubated at RT for 1.5 hours. (Sera were serially diluted into 1xPBS with a 10^0.5^ dilution factor (DF), starting from a dilution of 1:40. Monoclonal antibodies were serially diluted by a 10^0.5^ DF into 1xPBS. Starting concentrations for mAbs and sera are noted on ELISA graphs. The final well for each sample contained 1xPBS alone and guided the subtraction of the background signal calculation for each experiment.) Plates were washed 3x with PBS-T. For ELISAs that used murine sera as the source for the primary antibodies, the secondary antibody was rabbit pAb anti-mouse IgG-HRP (Abcam, AB97046). The secondary was added at 1:20,000 dilution (in PBS) on d34, d48, and d55, and it was added at 1:16000 on d13, d20, and d62. The secondary antibody solution was coated onto the plates for 1 hour at RT. For monoclonal antibodies, secondary goat pAb anti-human IgG-HRP (Abcam, AB97225) was added at 1:20,000 dilution (in PBS) for 1 hour at RT. Plates were washed three times with PBS-T and developed with 1-step ABTS Substrate Solution (ThermoFisher) for 45 minutes. 405nm absorbance values were measured using a plate reader. EC_50_ values were determined by non-linear regression (sigmoidal) using GraphPad Prism 10.2.3 software.

### Sera Depletion Assays

A/Japan/305/1957 (J’57), A/Bilthoven/21438/1971 (B’71), A/Darwin/6/2021 (D’21), and A/tern/Australia/G70C/1975 (Au’75) strains were cloned with N-terminus StrepII, i.e., TwinStrep, sequences, transiently expressed, and purified as described above. For the sera depletion assays, 65 columns of 500uL of Strep-Tactin^®^XT resin were prepared in 15mL gravity-flow columns and washed with PBS buffer. Based on prior studies related to antibody production kinetics^70^, excess StrepII tagged antigen was incubated with the Strep-Tactin resin for 1 hour, then eluted. Columns 1-35 were incubated with J’57 NA antigen. Columns 36-45 were incubated with B’71 NA antigen. Columns 46-55 were incubated with D’21 NA antigen. Columns 56-65 were incubated with Au’75 NA antigen. Separately, 555uL of PBS were mixed with 45µL of sera collected on d13 from each mouse for a 1/13.3 sera dilution. This was used as the stock of sera for each mouse for a given depletion experiment to minimize any experimental fluctuation due to pipetting. For mice m1-m35, 200uL of the diluted sera were incubated with the washed J’57-tagged Strep-Tactin resin columns 1-35, respectively. 200µL of m6-m15 sera mixtures were incubated with columns 36- 45, respectively. 200µL of m16-m25 sera mixtures were incubated with columns 46-55, respectively. Lastly, 200µL of m26-m55 sera mixtures were incubated with columns 56-65, respectively. After 2-4 hours at 4°C incubation, serum was eluted from each column and collected along with a 100µL of PBS column wash into the first rows of a 96-well PCR plate. The final concentration of the strain-depleted sera was therefore 1/20. These depleted sera were then serially diluted using a factor of 10^0.5^ for ELISA testing. Pre-depletion serum was taken from the same 1/13.3 diluted serum stock by taking 200µL of diluted serum and further diluting it with 100uL PBS for a final dilution of 1/20. The same was done for the sera that were collected on d55.

Pre-depletion serum and post J’57-depletion serum from m1-m5 were tested for reactivity against the J’57 strain (**figs. S9, S14**). The pre-depletion reactivity established the baseline J’57 reactivity value, and the post J’57 depletion reactivity was used as an internal control to verify successful J’57 depletion (*e.g.,* **fig. S13E**). Mice m1-m5 were also tested pre-depletion for reactivity to B’71, D’21, and Au’75 (**fig. S22**), respectively, to establish baseline levels of cross-reactivity in the control group. Mice m1-m5 were tested post J’57-depletion for any residual reactivity to B’71, D’21, and Au’75 (**figs. S9, S14**). Mice m6-m5 were tested pre-depletion for their reactivity to J’57 and B’71, post J’57-depletion for their reactivity to J’57 (internal control) and B’71 (*i.e.,* B’71 strain-specificity), and post B’71-depletion for their reactivity to J’57 (*i.e.,* J’57 strain-specificity) and B’71 (internal control). Analogous experiments were done for mice m16- m25 using J’57 and D’21 NAs, and for mice m26-m35 using J’57 and Au’75 NAs.

To minimize any variation in OD_405nm_ absorbance due to antigen coating, the same stock of 3µg/mL antigen concentration was used for all tested groups. Additionally, the 96-well (Corning) plates were organized to have all samples tested for reactivity against a given antigen on the same plates. For example, for the m6 serum that was tested against B’71, an entire plate was coated using the same B’71 3µg/mL diluted stock. Column 1 tested m6 pre-depletion serum vs. B’71, Column 2 tested m6 post-J’57 depletion vs. B’71, and Column 3 tested m6 post-B’71 depletion vs. B’71, etc. Since the number of tested samples for a given antigen exceeded one 96-well plate, the OD_405nm_ absorbance values were furthermore normalized for each antigen by dividing each OD_405nm_ data point by the average of the top 10 absorbance values for a given antigen.

### ELLA inhibition Assays

To test the ability of the sera to inhibit the NA activity of a given strain, tetramers of each strain were made, as NA tetramers have much greater catalytic activity than NA monomers^71,72^. J’57 NA was tetramerized using the tetrabrachion^73^ domain. B’71, D’21, and Au’75 NAs were tetramerized using VASP^74^ domains. (Choice of tetramerization domain was based solely on which domain helped the protein express better and more homogeneously.) Proteins were transiently expressed and purified as described above for the monomeric antigens. The EC_50_ of each NA activity was then determined. This was done through the following experiment setup: 100µL of 25µg/mL of fetuin (Sigma Aldrich, F3004) were coated onto 96- well plates and incubated overnight at 4°C. Plates were blocked with PBS, 1% BSA, 0.1% Tween-20 solutions and put on a shaker for 2 hours at RT. Plates were then washed 6x. Next, 100µL of recombinant NA_TB_, serially diluted 1:2 in DPBS with Ca/Mg (Gibco, Cat#: 14040117) with a starting concentration of 0.005µg/mL were added to the plate in columns 1-11. Column 12 was plated with DPBS alone. Plates were then placed in a 37°C incubator for 2 hours and subsequently washed with ELISA buffer 6x. Plates were incubated with 100µL of either 1 or 5µg/mL of HRP-conjugated lectin from Arachis hypogaea (peanut) (Sigma Aldrich, L7759) and incubated for 75-100min at RT on a plate shaker. Plates were then washed 6x and coated with 100uL of ABTS. After 45 min, plates were imaged. EC_50_ values of NA tetramers were determined by non-linear regression (sigmoidal) using GraphPad Prism 10.2.3 software. On d20, monomeric B’71 was used instead tetrameric B’71 due to poor tetramer expression, but the EC_50_ of the monomer was determined in the exact same way.

The ELLA inhibition assay was based on a standard protocol^55^. 100µL of 25µg/mL of fetuin (Sigma Aldrich, F3004) were coated onto 96-well plates and incubated overnight at 4°C. Plates were blocked with PBS, 1% BSA, 0.5% Tween-20 solutions and put on a shaker for 2 hours at RT. Plates were then washed 6x. A stock of 2x the EC_50_ of each strain was prepared and 65µL of solution was added to each well of a 96well PCR plate. Separately, 28µL of each tested mouse serum sample were added to 252µL of PBS with calcium, for a serum concentration of 1/10th. Serum was then serially diluted 1:2 down columns 1-11. Column 12 contained PBS with calcium. 65µL were aliquoted into the same 96 well plate as the 65µL of NA solution and pipetted 3x. After 5-10 minutes, 120µL of the sera/NA mixture were added to the fetuin-coated plate. [Sera from m1-m5 and a d0 sera negative control were tested against J’57, B’71, D’21, and Au’75. Sera from m6-m15 and a d0 sera negative control were tested against J’57 and B’71. Sera from m16-m25 and a d0 sera negative control were tested against J’57 and D’21. Finally, sera from m26-m35 and a d0 sera negative control were tested against J’57 and A’75.] Plates were placed into a 37°C for 90min, except for plates testing strain B’71, which incubated for 2hrs to give more time for NA activity to cleave the substrate. Plates were then washed 6x and coated with 100µL of HRP-lectin at 2µg/ml. They were then put on a RT shaker for 90 min. After washing the plates 6x, plates were coated with 150µL of ABTS. Plates were then imaged after exactly 45 min. Non-linear (sigmoidal) curves were generated for each serum sample using GraphPad Prism 10.2.3 software The loss of OD_405nm_ signal observed at the highest sera concentration (*i.e.,* dilution factor of 20) for the d0 negative control was used as the threshold for what would be considered ‘real’ signal due to specific sera antibody inhibition of NA rather than just the sera providing diffusion/viscosity inhibition of the NA activity. Sera from mice were then evaluated for the highest dilution factor at which the sera still outperformed the maximum inhibition of the d0 negative control. This was termed the sample’s MDFI, or the maximal dilution factor at which there was observable inhibition. Experimental group MDFIs were compared to the control. To avoid an observable hook-effect that impacted the regression analyses, all experiments for all samples were evaluated up until a dilution factor of 1280.

### Statistics

All analyses that compared data between only 2 groups of mice were performed using the unpaired, non-parametric Mann-Whitney test with an alpha value of 0.05. Analyses that compared the mean rank of one group to the mean rank of more than one group used the unpaired, non-parametric Kruskal-Wallis test with Dunn correction for multiple comparisons. The p-values cutoffs for significance were the following: 0.0332 (*), 0.0021 (**), 0.0002 (***), < 0.0001(****).

## ACKNOWLEDGMENTS

We thank Maria Ericsson and the Harvard Electron Microscopy Core for collection of negative stain images. We thank Daniel Maurer and Emerson Glassey for helpful discussions. We acknowledge support from NIGMS T32 GM0008313, NIH 1F30 AI181355 (to R.H.), NIH P01 AI089618 (A.G.S.), NIH AI155447, AI137057, AI153098 AI193280 (DL). This research has been funded in whole or part with federal funds under a contract from the National Institute of Allergy and Infectious Diseases, NIH contract 75N93019C00050 (A.G.S.).

## AUTHOR INFORMATION

### Author Contributions

R.H., A.G.S., and D.L. designed research; R.H. and F.A.N.M. performed research; R.H. and A.G.S. analyzed data; R.H. and A.G.S. wrote the paper. All authors commented on the manuscript.

### Correspondence and requests for materials should be addressed to

Aaron G. Schmidt

## References

1. Abduljaleel, Z. Decoding SARS-CoV-2 variants: Mutations, viral stability, and breakthroughs in vaccines and therapies. Biophys. Chem. 320–321, 107413 (2025).

2. Maurer, D. P., Vu, M. & Schmidt, A. G. Antigenic drift expands influenza viral escape pathways from recalled humoral immunity. Immunity 58, 716–727.e6 (2025).

3. Dingens, A. S., Arenz, D., Weight, H., Overbaugh, J. & Bloom, J. D. An Antigenic Atlas of HIV-1 Escape from Broadly Neutralizing Antibodies Distinguishes Functional and Structural Epitopes. Immunity 50, 520–532.e3 (2019).

4. Tenthorey, J. L., Emerman, M. & Malik, H. S. Evolutionary Landscapes of Host-Virus Arms Races. Annu. Rev. Immunol. 40, 271–294 (2022).

5. Caradonna, T. M. & Schmidt, A. G. Protein engineering strategies for rational immunogen design. Npj Vaccines 6, (2021).

6. Yassine, H. M. et al. Hemagglutinin-stem nanoparticles generate heterosubtypic influenza protection. Nat. Med. 21, 1065–1070 (2015).

7. Duan, H. et al. Glycan Masking Focuses Immune Responses to the HIV-1 CD4-Binding Site and Enhances Elicitation of VRC01-Class Precursor Antibodies. Immunity 49, 301–311 (2018).

8. Thornlow, D. N. et al. Altering the Immunogenicity of Hemagglutinin Immunogens by Hyperglycosylation and Disulfide Stabilization. Front. Immunol. 12, (2021).

9. Bajic, G. et al. Influenza Antigen Engineering Focuses Immune Responses to a Subdominant but Broadly Protective Viral Epitope. Cell Host Microbe 25, 827–835.e6 (2019).

10. Caradonna, T. M. et al. An Epitope-Enriched Immunogen Expands Responses to a Conserved Viral Site 1 2 3 4 Authors.

11. Doud, M. B., Hensley, S. E. & Bloom, J. D. Complete mapping of viral escape from neutralizing antibodies. PLOS Pathog. (2017) doi:10.1371/journal.ppat.1006271.

12. Lamson, D. T. et al. A modular platform to display multiple hemagglutinin subtypes on a single immunogen. eLife 13, (2025).

13. Arevalo, C. P. et al. A multivalent nucleoside-modified mRNA vaccine against all known influenza virus subtypes. Science 378, 899–904 (2022).

14. Kumari, M. et al. Multivalent mRNA Vaccine Elicits Broad Protection against SARS-CoV-2 Variants of Concern. Vaccines 12, 714 (2024).

15. Veneziano, R. et al. Role of nanoscale antigen organization on B-cell activation probed using DNA origami. Nat. Nanotechnol. 15, 716–723 (2020).

16. Slifka, M. K. & Amanna, I. J. Role of Multivalency and Antigenic Threshold in Generating Protective Antibody Responses. Front. Immunol. 10, (2019).

17. Kelly, H. G., Kent, Stephen J & and Wheatley, A. K. Immunological basis for enhanced immunity of nanoparticle vaccines. Expert Rev. Vaccines 18, 269–280 (2019).

18. Li, Y. et al. Enhancing immunogenicity and transmission-blocking activity of malaria vaccines by fusing Pfs25 to IMX313 multimerization technology. Sci. Rep. 6, 18848 (2016).

19. Cui, X. et al. Immunization of Rabbits with Recombinant Human Cytomegalovirus Trimeric versus Monomeric gH/gL Protein Elicits Markedly Higher Titers of Antibody and Neutralization Activity. Int. J. Mol. Sci. 20, 3158 (2019).

20. Tsai, S. J., Black, S. K. & Jewell, C. M. Leveraging the modularity of biomaterial carriers to tune immune responses. Adv. Funct. Mater. 30, 2004119 (2020).

21. Brouwer, P. J. M., et al. Immunofocusing and enhancing autologous Tier-2 HIV-1 neutralization by displaying Env trimers on two-component protein nanoparticles. Npj Vaccines 6, (2021).

22. Ols, S. et al. Multivalent antigen display on nanoparticle immunogens increases B cell clonotype diversity and neutralization breadth to pneumoviruses. Immunity 56, 2425–2441.e14 (2023).

23. Kato, Y. et al. Multifaceted Effects of Antigen Valency on B Cell Response Composition and Differentiation In Vivo. Immunity 53, 548–563.e8 (2020).

24. Li, M. et al. Enhancing humoral and mucosal immune response of PED vaccine candidate by fusing S1 protein to nanoparticle multimerization. Vet. Microbiol. 290, 110003 (2024).

25. Abbott, R. K. et al. Precursor Frequency and Affinity Determine B Cell Competitive Fitness in Germinal Centers, Tested with Germline-Targeting HIV Vaccine Immunogens. Immunity 48, 133–146.e6 (2018).

26. Mohsen, M. O. & Bachmann, M. F. Virus-like particle vaccinology, from bench to bedside. Cell. Mol. Immunol. 19, 993–1011 (2022).

27. King, N. P. et al. Accurate design of co-assembling multi-component protein nanomaterials. Nature 510, 103–108 (2014).

28. Cohen, A. A. et al. Mosaic nanoparticles elicit cross-reactive immune responses to zoonotic coronaviruses in mice. Science 735–741 (2021).

29. Kanekiyo, M. et al. Mosaic nanoparticle display of diverse influenza virus hemagglutinins elicits broad B cell responses. Nat. Immunol. 20, 362–372 (2019).

30. Brinkkemper, M. et al. Mosaic and mixed HIV-1 glycoprotein nanoparticles elicit antibody responses to broadly neutralizing epitopes. PLOS Pathog. 20, e1012558 (2024).

31. Boyoglu-Barnum, S. et al. Quadrivalent influenza nanoparticle vaccines induce broad protection. Nature 592, 623–628 (2021).

32. Dosey, A. et al. Combinatorial immune refocusing within the influenza hemagglutinin RBD improves cross-neutralizing antibody responses. Cell Rep. 42, 113553 (2023).

33. Pascha, M. N. et al. Nanoparticle display of neuraminidase elicits enhanced antibody responses and protection against influenza A virus challenge. Npj Vaccines 9, 1–14 (2024).

34. Allen, J. D. & Ross, T. M. Bivalent H1 and H3 COBRA Recombinant Hemagglutinin Vaccines Elicit Seroprotective Antibodies against H1N1 and H3N2 Influenza Viruses from 2009 to 2019. J. Virol. 96, e01652–21 (2022).

35. McAuley, J. L., Gilbertson, B. P., Trifkovic, S., Brown, L. E. & McKimm-Breschkin, J. L. Influenza virus neuraminidase structure and functions. Front. Microbiol. 10, (2019).

36. Carter, T. & Iqbal, M. The Influenza A Virus Replication Cycle: A Comprehensive Review. Viruses 16, 316 (2024).

37. Liu, C., Eichelberger, M. C., Compans, R. W. & Air, G. M. Influenza type A virus neuraminidase does not play a role in viral entry, replication, assembly, or budding. J. Virol. 69, 1099–1106 (1995).

38. Krammer, F. et al. NAction! how can neuraminidase-based immunity contribute to better influenza virus vaccines? mBio 9, (2018).

39. Creytens, S., Pascha, M. N., Ballegeer, M., Saelens, X. & de Haan, C. A. M. Influenza Neuraminidase Characteristics and Potential as a Vaccine Target. Front. Immunol. 12, (2021).

40. Eichelberger, M. C. & Monto, A. S. Neuraminidase, the Forgotten Surface Antigen, Emerges as an Influenza Vaccine Target for Broadened Protection. J. Infect. Dis. 219, S75–S80 (2019).

41. Zhang, X. & Ross, T. M. Anti-neuraminidase immunity in the combat against influenza. Expert Rev. Vaccines 23, 474–484 (2024).

42. Stadlbauer, D. et al. Broadly protective human antibodies that target the active site of influenza virus neuraminidase.

43. Momont, C. et al. A pan-influenza antibody inhibiting neuraminidase via receptor mimicry. Nature (2023) doi:10.1038/s41586-023-06136-y.

44. Yasuhara, A. et al. A broadly protective human monoclonal antibody targeting the sialidase activity of influenza A and B virus neuraminidases. Nat. Commun. 13, 6602–6602 (2022).

45. Johansson, B. E., Matthews, J. T. & Kilbourne, E. D. Supplementation of conventional influenza A vaccine with purified viral neuraminidase results in a balanced and broadened immune response. Vaccine 16, 1009–1015 (1998).

46. Monto, A. S. et al. Antibody to Influenza Virus Neuraminidase: An Independent Correlate of Protection. J. Infect. Dis. 212, 1191–1199 (2015).

47. Zakeri, B. et al. Peptide tag forming a rapid covalent bond to a protein, through engineering a bacterial adhesin. doi:10.1073/pnas.1115485109/-/DCSupplemental.

48. Klein, J. S. et al. Design and characterization of structured protein linkers with differing flexibilities. in vol. 27 325–330 (Oxford University Press, 2014).

49. Cho, K. J. et al. The Crystal Structure of Ferritin from *Helicobacter pylori* Reveals Unusual Conformational Changes for Iron Uptake. J. Mol. Biol. 390, 83–98 (2009).

50. Zhu, X. et al. Structural Basis of Protection against H7N9 Influenza Virus by Human Anti-N9 Neuraminidase Antibodies. Cell Host Microbe 26, 729–738.e4 (2019).

51. Lei, R. et al. Leveraging vaccination-induced protective antibodies to define conserved epitopes on influenza N2 neuraminidase. Immunity 56, 2621–2634.e6 (2023).

52. Lederhofer, J. et al. Protective human monoclonal antibodies target conserved sites of vulnerability on the underside of influenza virus neuraminidase. Immunity 57, 574–586.e7 (2024).

53. Harris, L. J., Larson, S. B., Hasel, K. W. & McPherson, A. Refined Structure of an Intact IgG2a Monoclonal Antibody,. Biochemistry 36, 1581–1597 (1997).

54. Schmidt, A. G. et al. Viral receptor-binding site antibodies with diverse germline origins. Cell 161, 1026–1034 (2015).

55. Gao, J., Couzens, L. & Eichelberger, M. C. Measuring Influenza Neuraminidase Inhibition Antibody Titers by Enzyme-linked Lectin Assay. J. Vis. Exp. JoVE 54573 (2016) doi:10.3791/54573.

56. Leang, S. K. & Hurt, A. C. Fluorescence-based neuraminidase inhibition assay to assess the susceptibility of influenza viruses to the neuraminidase inhibitor class of antivirals. J. Vis. Exp. (2017) doi:10.3791/55570.

57. Lainšček, D. et al. A Nanoscaffolded Spike-RBD Vaccine Provides Protection against SARS-CoV- 2 with Minimal Anti-Scaffold Response. Vaccines 9, 431 (2021).

58. Mesin, L., Ersching, J. & Victora, G. D. Germinal Center B Cell Dynamics. Immunity 45, 471–482 (2016).

59. Cyster, J. G. & Allen, C. D. C. B Cell Responses: Cell Interaction Dynamics and Decisions. Cell 177, 524–540 (2019).

60. Klein, F. et al. Somatic mutations of the immunoglobulin framework are generally required for broad and potent HIV-1 neutralization. Cell 153, 126–138 (2013).

61. Pilewski, K. A. et al. Functional HIV-1/HCV cross-reactive antibodies isolated from a chronically co-infected donor. Cell Rep. 42, 112044 (2023).

62. Wang, M. et al. Rapid clonal expansion and somatic hypermutation contribute to the fate of SARS-CoV-2 broadly neutralizing antibodies. J. Immunol. 214, 278–289 (2025).

63. Walz, L., Kays, S.-K., Zimmer, G. & von Messling, V. Neuraminidase-Inhibiting Antibody Titers Correlate with Protection from Heterologous Influenza Virus Strains of the Same Neuraminidase Subtype. J. Virol. 92, e01006–18 (2018).

64. Wohlbold, T. J. et al. Vaccination with adjuvanted recombinant neuraminidase induces broad heterologous, but not heterosubtypic, cross-protection against influenza virus infection in mice. mBio 6, e02556 (2015).

65. Kaplan, B. S. et al. A neuraminidase-based inactivated influenza virus vaccine significantly reduced virus replication and pathology following homologous challenge in swine. Vaccine 46, 126574 (2025).

66. Stadlbauer, D. et al. Antibodies targeting the neuraminidase active site inhibit influenza H3N2 viruses with an S245N glycosylation site. Nat. Commun. 13, 7864 (2022).

67. Schiepers, A. et al. Molecular fate-mapping of serum antibody responses to repeat immunization. Nature (2023) doi:10.1038/s41586-023-05715-3.

68. Gostic, K. M. et al. Childhood immune imprinting to influenza A shapes birth year-specific risk during seasonal H1N1 and H3N2 epidemics. PLoS Pathog. 15, e1008109 (2019).

69. Pingoud, A., Fuxreiter, M., Pingoud, V. & Wende, W. Type II restriction endonucleases: structure and mechanism. Cell. Mol. Life Sci. CMLS 62, 685–707 (2005).

70. Hauge, S., Madhun, A., Cox, R. J. & Haaheim, L. R. Quality and Kinetics of the Antibody Response in Mice after Three Different Low-Dose Influenza Virus Vaccination Strategies. Clin. Vaccine Immunol. CVI 14, 978–983 (2007).

71. Zanin, M. et al. An Amino Acid in the Stalk Domain of N1 Neuraminidase Is Critical for Enzymatic Activity. J. Virol. 91, 10.1128/jvi.00868-16 (2017).

72. Deroo, T., Min Jou, W. & Fiers, W. Recombinant neuraminidase vaccine protects against lethal influenza. Vaccine 14, 561–569 (1996).

73. Streltsov, V. A., Schmidta, P. M. & Breschkina, J. L. M. K. Structure of an influenza a virus n9 neuraminidase with a tetrabrachion-domain stalk. Acta Crystallogr. Sect. F Struct. Biol. Commun. 75, 89– 97 (2019).

74. Kü, K. et al. The VASP tetramerization domain is a right-handed coiled coil based on a 15-residue repeat. PNAS 101, 17027–17032 (2004).

