## Supplemental Files for "Neuraminidase-on-a-string nanoparticles probe how antigenic distance shapes elicited humoral immunity"

**A** Ferritin nanoparticle (NP) protomer secreted amino acid sequence:

AVDTLSGLSSEQGQSGDMTIEEDSATHIKFSK
RDEDGKELAGATMELRDSSGKTISTWISDGQ
VKDFYLYPGKYTFVETAAPDGYEVATAITFTVN
EQGQVTVNGKATKGDAHIGGGSGGGSHHHH
HHHHGGGSGGGSGAGSSGSSGSSGSS**DIIK**
LLNEQVNKEMNSSNLYMSMSSWCYTHSLDG
AGLFLFDHAAEEYEHAKKLIVFLNHHNVPVQL
TSISAPEHKFEGLTQIFQKAYEHEQHISESINNI
VDHAIKGKDHHTFNFLQWYVSEQHEEEVLFK
DILDKIELIGNEHHGLYHADQYVKGIAKSRKS

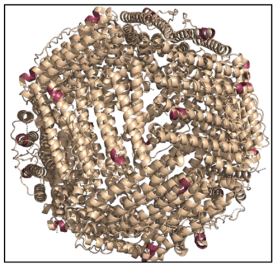

**B**

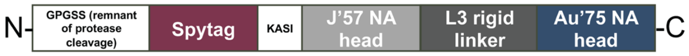


GPGSS-**MAHIVMVDAYKPTKGSG**-GA-J'57head-
**TGSGPVPSTPPTPSPSTPPTPSPSGGSGNSSGSGGSPVPSTPPTPS**
**PSTPPTPSPSAS**-Au'75head

**C**

```
Consensus                        XVEYR NWSKP QCXIT GFAPF SKDNS IRLSA GGDIW VTREP YVSCD PXKCY   50
A/Japan/305/1957 (H2N2)          V.... ..... ..Q.. ..... ..... ..... ..... ..... ..... .G...   50
A/Bilthoven/21438/1971 (H3N2)    L.... ..... ..K.. ..... ..... ..... ..... ..... ..... .G...   50
A/Darwin/6/2021 (H3N2)           -A... ..... ..G.. ..... ..... ..... ..... ..... ..... LD...   49
A/tern/Australia/G70C/1975 (H11N9) ---FN .LT.G L.T.N SWHIY G...A V.IGE DS.VL ..... ..... .DE.R   47

Consensus                        QFALG QGTTL DNKHS NGTIH DRIPX RTLLM NELGV PFHLG TKQV- CIAWS   99
A/Japan/305/1957 (H2N2)          ..... ..... ..... ..... ....H ..... ..... ..... ....- .V...   99
A/Bilthoven/21438/1971 (H3N2)    ..... ..... ..... .D... ....H ..... ..... ..... .R..- .....   99
A/Darwin/6/2021 (H3N2)           ..... ..... N.V.. .N.VR ..T.Y ..... ..... ..... ....- .....   98
A/tern/Australia/G70C/1975 (H11N9) FY..S ....I RG... ..... ..SQY .A.IS WP.SS .PTVY NSR.E ..G..   97

Consensus                        SSSCH DGKAW LHVCX TGDDK NATAS FIYBG RLVDS IGSWS QNILR TQESE  149
A/Japan/305/1957 (H2N2)          ..... ..... ....V ....R ..... ...D. ..... ..... ..... .....  149
A/Bilthoven/21438/1971 (H3N2)    ..... ..... ....V ..... ..... ...D. ..... ..... ..... .....  149
A/Darwin/6/2021 (H3N2)           ..... ..... ....I ..... ..... ...N. ..... VV... ND... .....  148
A/tern/Australia/G70C/1975 (H11N9) .T... ...TR MSI.I S.PNN ..S.V IW.NR .P.TE .NT.A R.XX. .....  147

Consensus                        CVCIN GTCTV VMTDG SAXGK ADTRI LFIXE GKIVH ISPLS GSAQH XEECS  199
A/Japan/305/1957 (H2N2)          ..... ..... ..... ..S.R ..... ...K. ..... ..... ..... I....  199
A/Bilthoven/21438/1971 (H3N2)    ..... ..... ..... ..S.. ..... ...E. ..... ..... ..... V....  199
A/Darwin/6/2021 (H3N2)           ..... ..... ..... N.T.. ...K. ...E. ..... T.K.. ..... V....  198
A/tern/Australia/G70C/1975 (H11N9) ...H. .V.P. .F... ..T.P .E... YYFK. ...LK WE..A .T.K. I....  197

Consensus                        CYPRY PGVRC ICRDN WKGSN RPVID INXKD YSIDS SYVCS GLVGD TPRNN  249
A/Japan/305/1957 (H2N2)          ..... .D... ..... ..... ..... ..ME. ..... ..... ..... ....D  249
A/Bilthoven/21438/1971 (H3N2)    ..... ..... ..... ..... ...V. ..V.. ..... ..... ..... .....  249
A/Darwin/6/2021 (H3N2)           ..... ..... V.... ..... ..I.. ..I.. H..V. R.... ..... ...KS  248
A/tern/Australia/G70C/1975 (H11N9) ..GER AEIT. T.... .Q... ....R .DPVA MTHT. Q.I.. PVLT. N..P.  247

Consensus                        DSSSS SXCRB PNNER GNHGV KGWAF DDGND VWMGR TISKD SRSGY ETFKV  299
A/Japan/305/1957 (H2N2)          ....N .N..D ..... ..P.. ..... .N.D. ..... ..... ..... .....  299
A/Bilthoven/21438/1971 (H3N2)    .R... .Y..N ..... ..... ..... ..... ..... ..... ..... .....  299
A/Darwin/6/2021 (H3N2)           ..... .H.LN ....K .D... ..... ..... ..... ..NET ..L.. .....  298
A/tern/Australia/G70C/1975 (H11N9) .PTVG K-.ND .YPGN N.N.. ..FSY L..VN T.L.. ...IA ..... .ML..  296

Consensus                        IGGWS TPXSK SQINR QVIVD NXBXS GYSGI FS--- VEGKS CINRC FYVEL  346
A/Japan/305/1957 (H2N2)          ..... ..N.. ..V.. ..... .NNW. ..... ..--- ..... ..... .....  346
A/Bilthoven/21438/1971 (H3N2)    ..... ..N.. ..... ..... SDNR. ..... ..--- ..... ..... .....  346
A/Darwin/6/2021 (H3N2)           VE... N.K.. L.... ..... RGDR. ..... ..--- ..... ..... .....  345
A/tern/Australia/G70C/1975 (H11N9) PNALT DDK.. -PTQG .T..L .TDW. ....S .MDYW A..E- .YRA. .....  344

Consensus                        IRGRP QETRV WWTSN SIVVF CGTSG TYGTG SWPDG ANINL MPI          389
A/Japan/305/1957 (H2N2)          ..... ..... ..... ..... ..... ..... ..... ....F ...          389
A/Bilthoven/21438/1971 (H3N2)    ....E ..... ..... ..... ..... ..... ..... .D... ...          389
A/Darwin/6/2021 (H3N2)           ....K E..E. L.... ..... ..... ..... ..... ..LS. .H.          388
A/tern/Australia/G70C/1975 (H11N9) ..... K.DK. ..... ...SM .SSTE FL.QW D.... .K.EY FL-          386
```

**fig. S1: Neuraminidase-on-a-string (NoaS) nanoparticle (NP) components. (A)** Amino acid sequence for the *Helicobacter Pylori* ferritin protomer. The ribbon diagram for the ferritin nanoparticle (NP) is shown in gold, with the N-terminus of each protomer highlighted in raspberry (PDB 3BVE[1]). **(B)** Schematic of the NoaS design. The N-terminus of the dimer protein began with a GSG remnant from an HRV 3C cleavage site, followed by the amino acid sequence for the SpyTag[2] site (listed below in raspberry letters), then the amino acids 'GA,' which encode the KasI enzyme cloning site. The first NA head was immediately followed by the L3-rigid protein linker[3] and then followed by the amino acids for the second NA head to form a SpyTag, rigidly-linked NoaS dimer. The example here shows the schematic for the J'57-Linker-Au'75 NoaS, which was the immunogen used for group 7. **(C)** Amino acid alignment of the four NA heads used in the NoaS NP groups: A/Japan/305/1957, A/Bilthoven/21438/1971, A/Darwin/6/2021, and A/tern/Australia/G70C/1975.

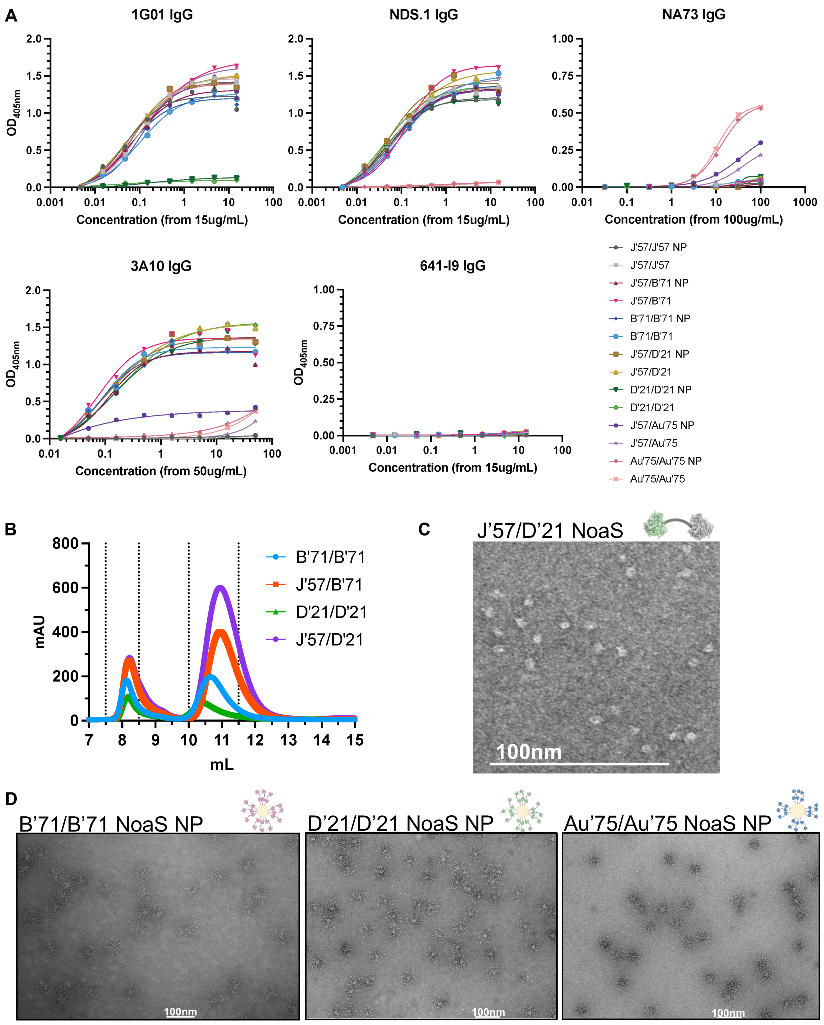


**fig. S2: Biochemical characterization of NoaS NP immunogens. (A)** ELISA binding curves of mAbs used to confirm the accessibility and structural integrity of the NA heads on the NoaS NPs. 1G01[4] mAb binds J'57, B'71, and Au'75 NAs and does not bind D'21 NA. NDS.1[5] mAb binds all three N2s and does not bind Au'75 NA. NA73[6] mAb has partial albeit selective reactivity to the Au'75 N9 NA. 3A10[7] mAb does not bind the J'57 NA and was used to selectively verify the structural integrity of the B'71 and D'21 NAs. **(B)** Size exclusion chromatograms for the group 2, 3, 4, and 5 NoaS NPs. **(C)** Negative stain electron microscopy images of the group 2, 4, and 6 NoaS NP immunogens.

A

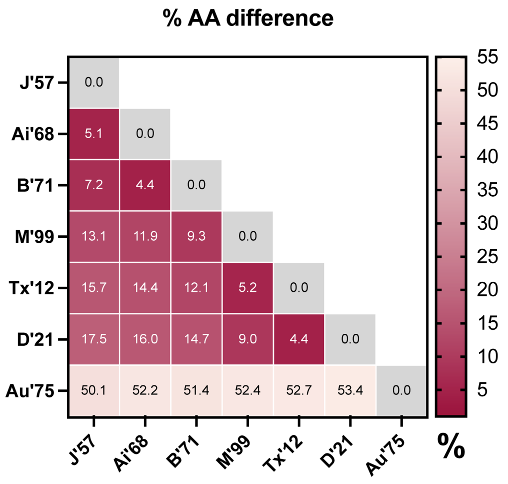


B

| Strain: | Abridged: | # of PNGs: |
|---|---|---|
| A/Japan/305/1957 | J'57 | 5 |
| A/Aichi/2/1968 | Ai'68 | 5 |
| A/Bilthoven/21438/1971 | B'71 | 5 |
| A/Moscow/10/1999 | M'99 | 7 |
| A/Texas/50/2012 | Tx'12 | 6 |
| A/Darwin/6/2021 | D'21 | 7 |
| A/tern/Australia/G70C/1975 | Au'75 | 3 |

**fig. S3: Historical NA panel used to assess elicited NA-breadth in ELISA assays. (A)** Heatmap of the percentage differences in amino acid composition within the NA heads for the seven NA strains included in the breadth panel. **(B)** Complete names of the seven strains, their abbreviations used, and the number of predicted N-linked glycosylation sites (PNGs) within each NA head.

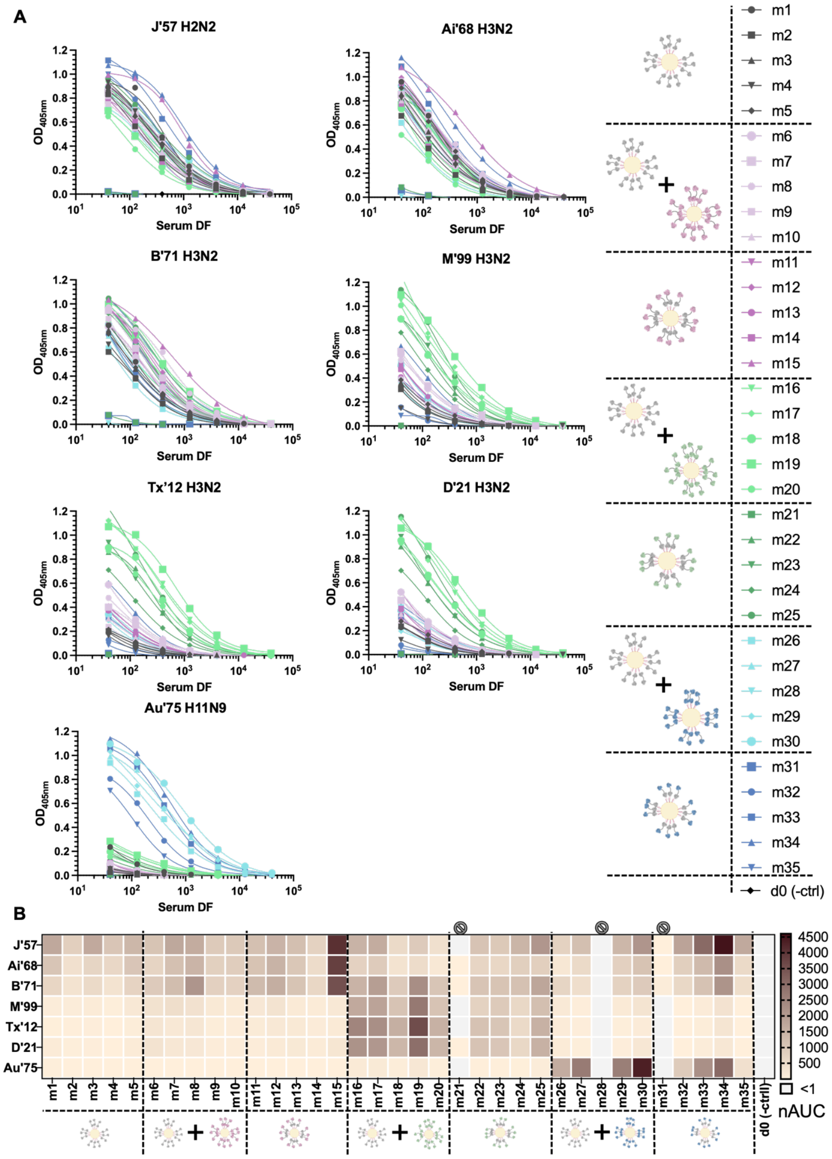


**fig. S4: Reactivity of sera from d20 to historical NA panel. (A)** ELISA data of the d20 sera from each mouse, tested against the panel of historical NAs. Curves represent the $OD_{405nm}$ absorbance values for each serial dilution of sera, normalized per antigen. **(B)** The same ELISA data shown as a heatmap of normalized area under the curve (AUC) values. Three mice, m21, m28, and m31, did not respond to the immunizations, and their data were excluded from any statistical analyses.

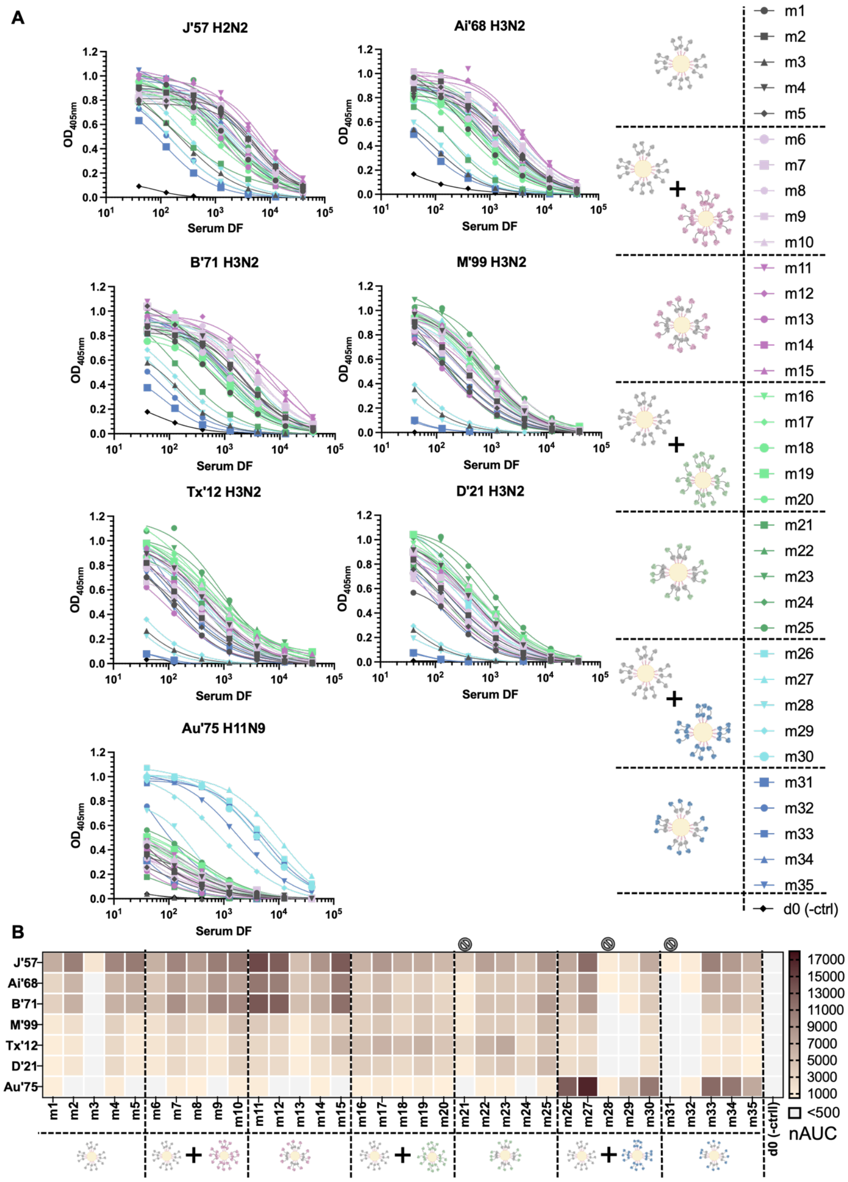


**fig. S5: Reactivity of sera from d48 to historical NA panel. (A)** ELISA reactivity of d48 sera from each mouse, tested against the panel of historical NA antigens. Data shown is the $OD_{405nm}$ absorbance values for each serial dilution of sera, normalized per antigen. **(B)** The same ELISA data represented as a heatmap of normalized area under the curve (AUC) values. Three mice, m21, m28, and m31, did not respond to the prime immunizations, and their data were therefore excluded from any statistical analyses.

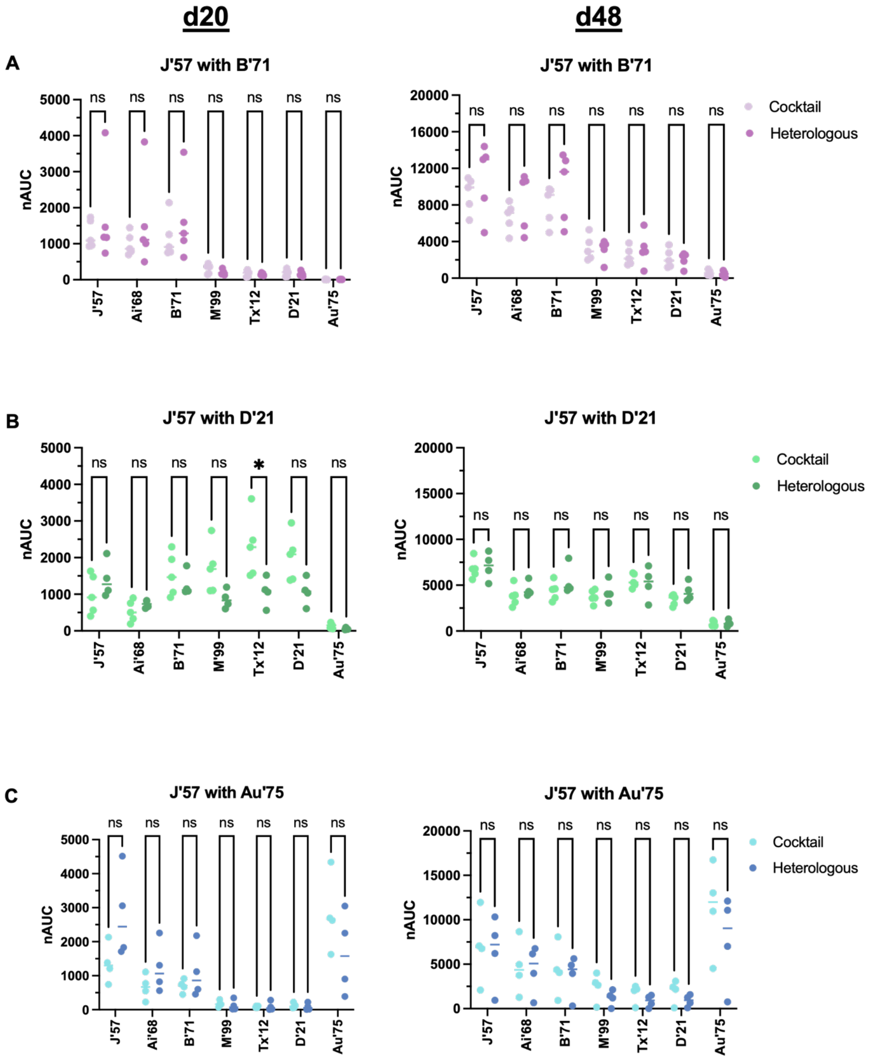


**fig. S6: Reactivities of mouse sera against each antigen in the NA panel. (A)** Groups 2 and 3 (J'57 NA paired with B'71 NA). **(B)** Groups 4 and 5 (J'57 NA paired with D'21 NA). **(C)** Groups 6 and 7 (J'57 NA paired with Au'75 NA). Groups 2, 4, and 6 were mice that received a cocktail of two homologous NoaS NPs. Groups 3, 5, and 7 were mice that received an equimolar heterologous NoaS NP immunization incorporating the same two strains. Statistical significance is noted only for groups 4 and 5 reactivity against Tx'12 NA (p-value: *0.03). The data from groups 2 and 3, 4 and 5, and 6 and 7 were pooled to reflect the differences in reactivity profiles that varied based on which strains were included in the immunizations. Statistical significance was determined using Mann-Whitney tests.

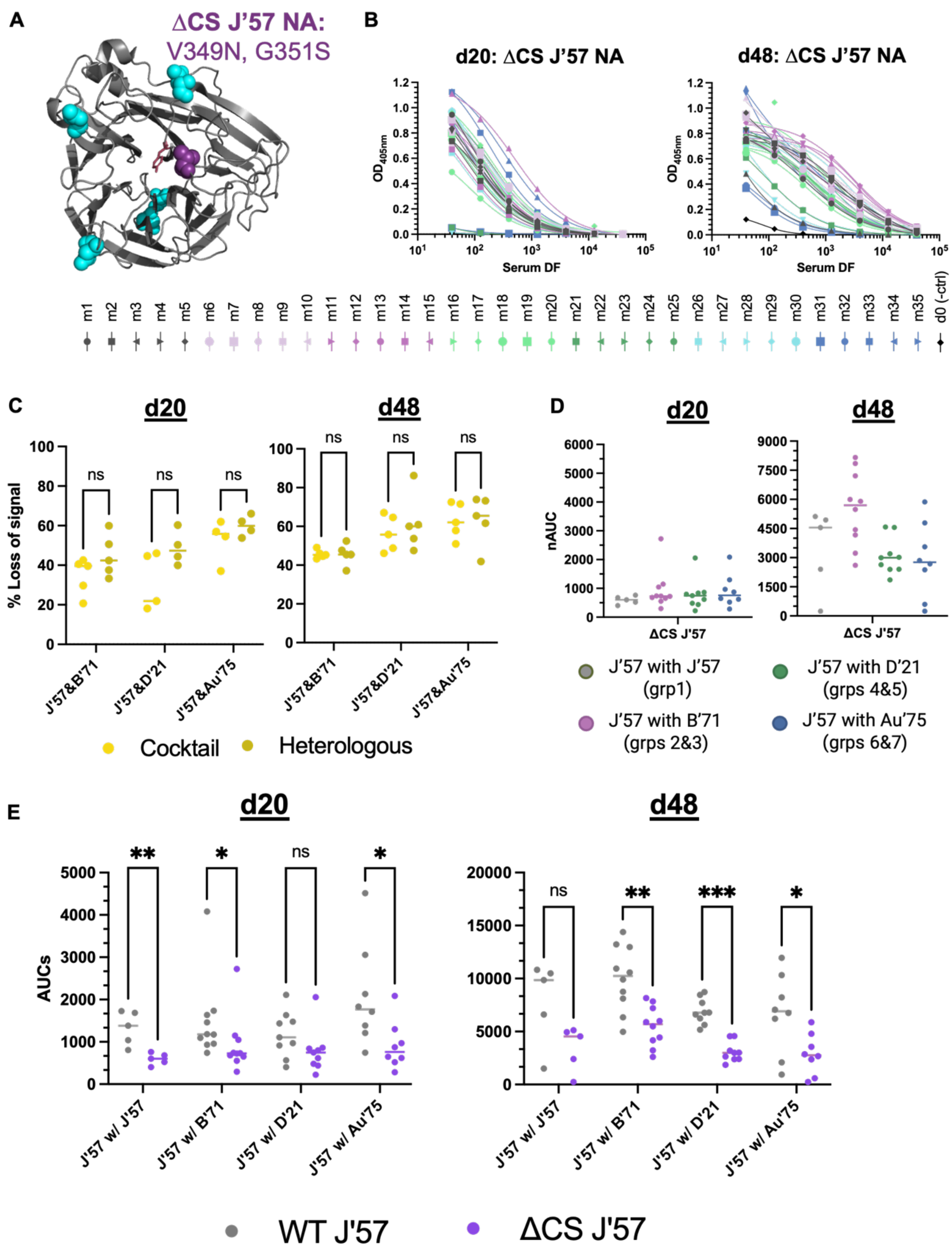


**fig. S7: Assessment of sera focusing onto J'57 catalytic site (CS). (A)** Cartoon representation of the monomeric J'57 NA head (PDB 6Q20[4]) in gray, showing the locations of its 5 wildtype PNG sites in cyan and the engineered, CS-blocking PNG site (V349N, G351S) in purple. **(B)** Normalized ELISA reactivity curves of d20 and d48 sera tested against the ΔCS-J'57 NA head. **(C)** Loss of reactivity between the wildtype J'57 NA head and the ΔCS-J'57 was assessed for each mouse serum sample, broadly reflecting the J'57-reactivity within a given serum sample that was directed toward the CS of J'57 (referenced as the % loss of signal). No statistically significant differences in the % loss of signal for the ΔCS-J'57 antigen in the cocktail vs. heterologous groups (*i.e.,* groups 2, 4, 6 vs. groups 3, 5, 7) were observed. Data from groups 2 and 3 (J'57 with B'71), 4 and 5 (J'57 with D'21), and 6 and 7 (J'57 with Au'75) were pooled to reflect the differences in reactivity that varied between groups based on which strains were included in their immunizations. **(D)** ELISA reactivity for the ΔCS-J'57 antigen on d20 and d48. No statistically significant differences in reactivity were observed across groups. **(E)** Assessing ELISA AUCs of wildtype J'57 NA compared to ΔCS-J'57 NA. The degree of AUC signal lost against ΔCS-J'57 varied across the groups based on what strain J'57 was paired with in an immunization. Statistically significant loss of signal was observed for the J'57/J'57 control group on d20 (p-value: **0.008) but not d48. On both d20 and d48, significance of signal loss was noted for the J'57/B'71 groups (p-values: *0.01 and **0.002, respectively). On d20, loss of signal for J'57/D'21 was not statistically significant, but significance was noted by d48 (p-value: ***<0.0001). Statistical significance of signal loss was noted for groups that received J'57/Au'75 on both d20 and d48 (p-values: *0.01 and *0.04, respectively).

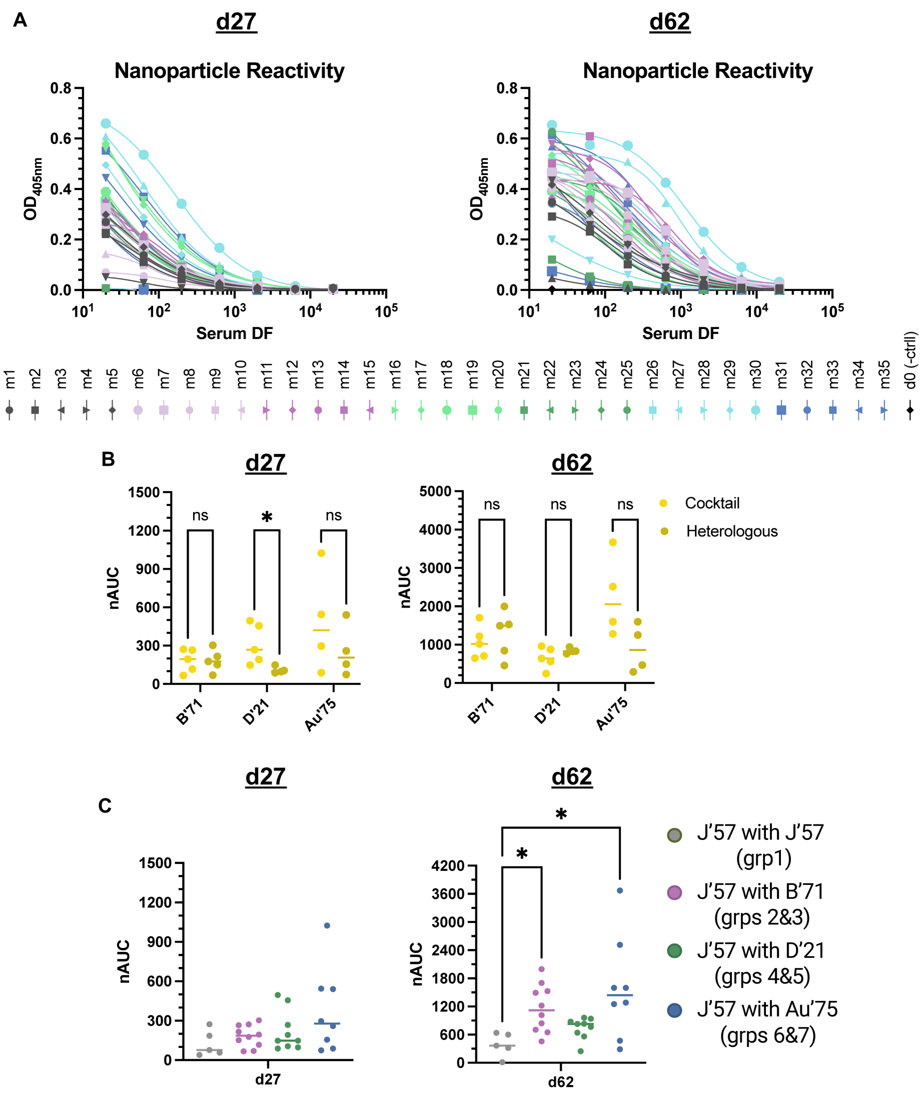


**fig. S8: Reactivity of the sera to the naked ferritin nanoparticle. (A)** ELISA curves of all mice from pre-boost d27 and post-boost d62 to the naked ferritin NP. **(B)** d27 and d62 sera reactivity against the naked ferritin NP, shown as reactivities of homologous cocktail groups versus their heterologous NoaS NP groups. Except for group 4 versus group 5 on d27 (p-value: *0.03), there were no statistically significant differences in reactivity to the naked NP observed. Data for each pair of strains were therefore evaluated collectively. **(C)** Reactivity to the naked NP was comparable across group pairs on d27. On d62, significant differences were observed for the naked NP reactivity between the control group and the groups that received B'71 NA (p-value: *0.0323) and Au'75 NA (p-value: *0.0159) as their second NA component. Statistical significance was evaluated using Mann-Whitney.

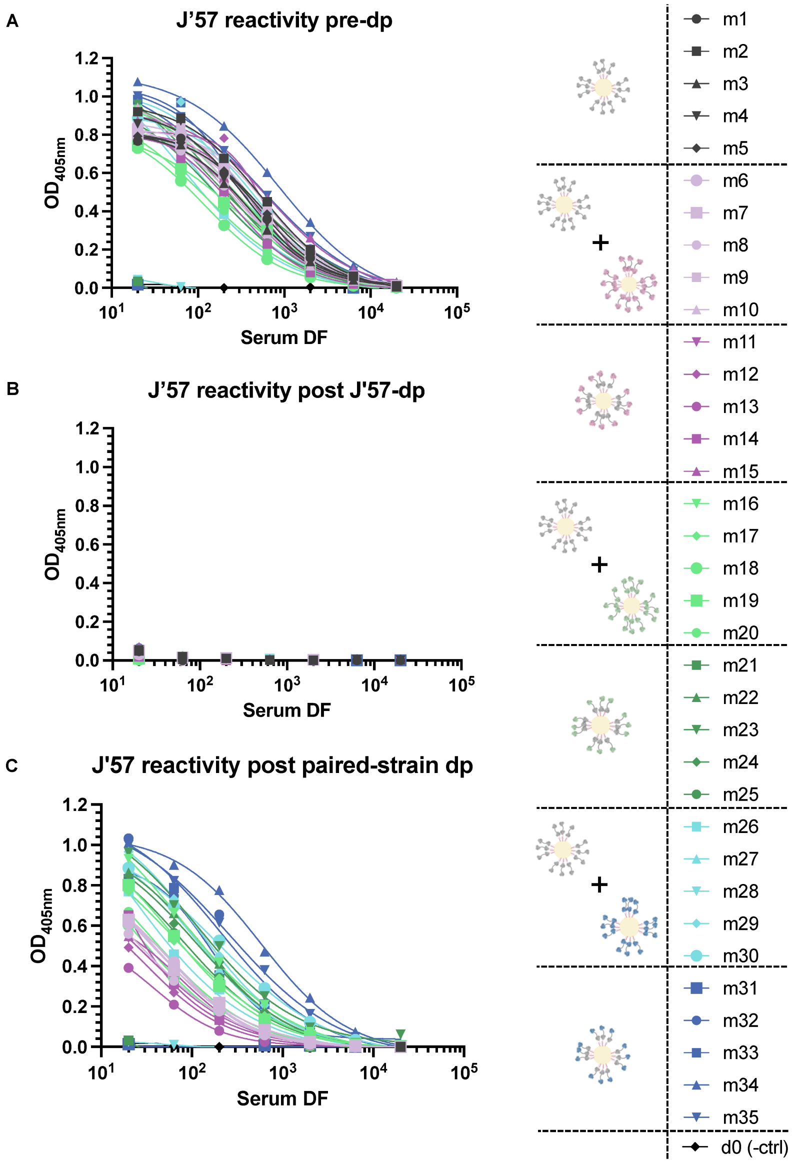


**fig. S9: Day 13 sera depletion studies for the J’57 NA strain. (A)** ELISA reactivity of the sera from all immunized mice to J’57 NA prior to any NA depletions. Sera samples from mice m1-m35 were depleted with J’57 NA. J’57 NA-depleted sera samples were divided into two equal parts. One part was tested against J’57 NA, as shown in **(B)**, verifying that there was no remaining J’57 NA reactivity in the sera. The other part of the J’57 NA-depleted sera was tested against the remaining strains. **(C)** Sera from mice that were depleted with B’71 NA (m6-m15), D’21 NA (m16-m25), and Au’75 NA (m26-m35) were tested for remaining reactivity against J’57 NA.

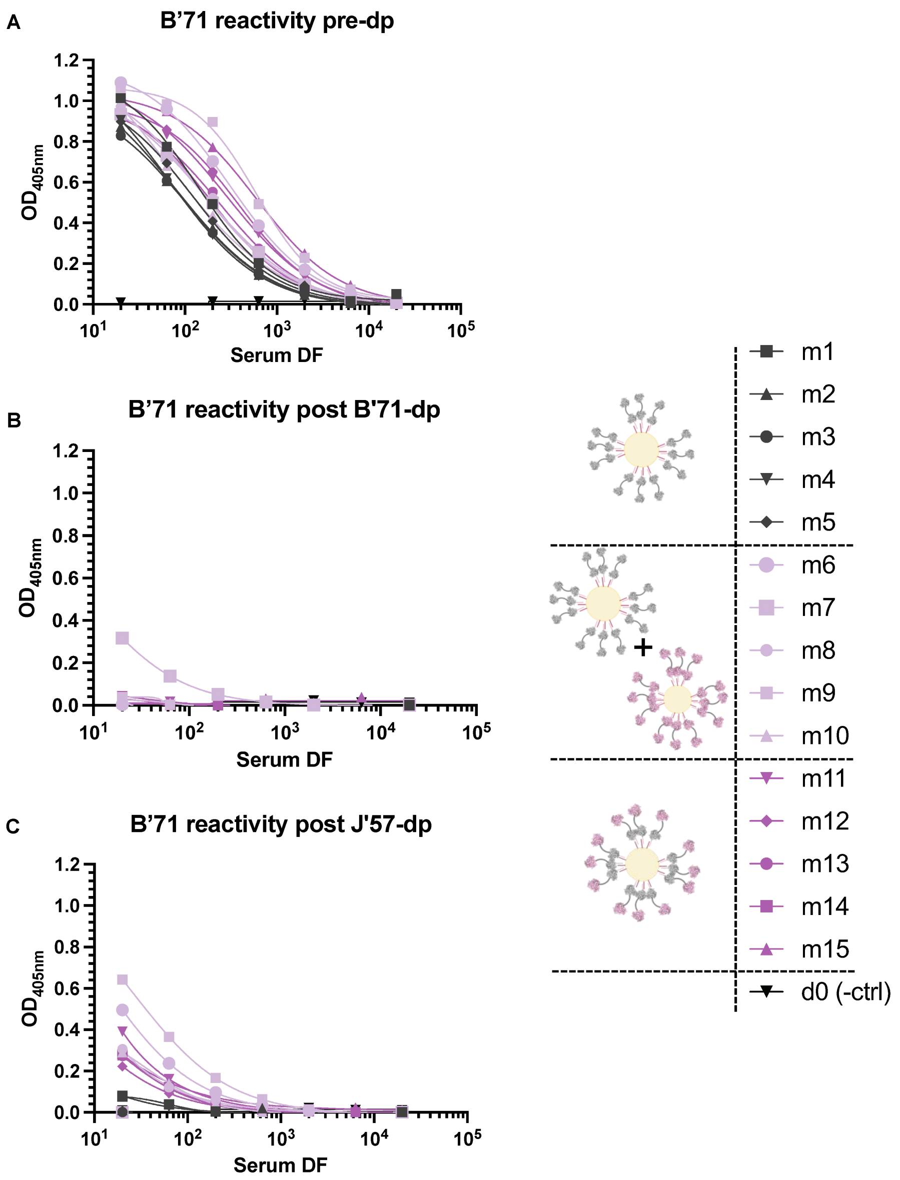


**fig. S10: Day 13 sera depletion studies for the B'71 NA. (A)** ELISA reactivity of the sera from mice m1-m15 to B'71 NA prior to any strain depletions. Sera samples for mice m1-m15 were depleted with B'71 NA antigen for any B'71 reactivity. B'71 NA-depleted sera samples were divided into two equal parts. One part was tested against B'71 NA, as shown in **(B)**, verifying that there was no remaining B'71 NA reactivity in the sera. (The other part of the B'71 NA-depleted sera were tested against J'57 NA.) **(C)** Sera from mice that were depleted for J'57 NA (m1-m15) were tested for remaining reactivity against B'71 NA.

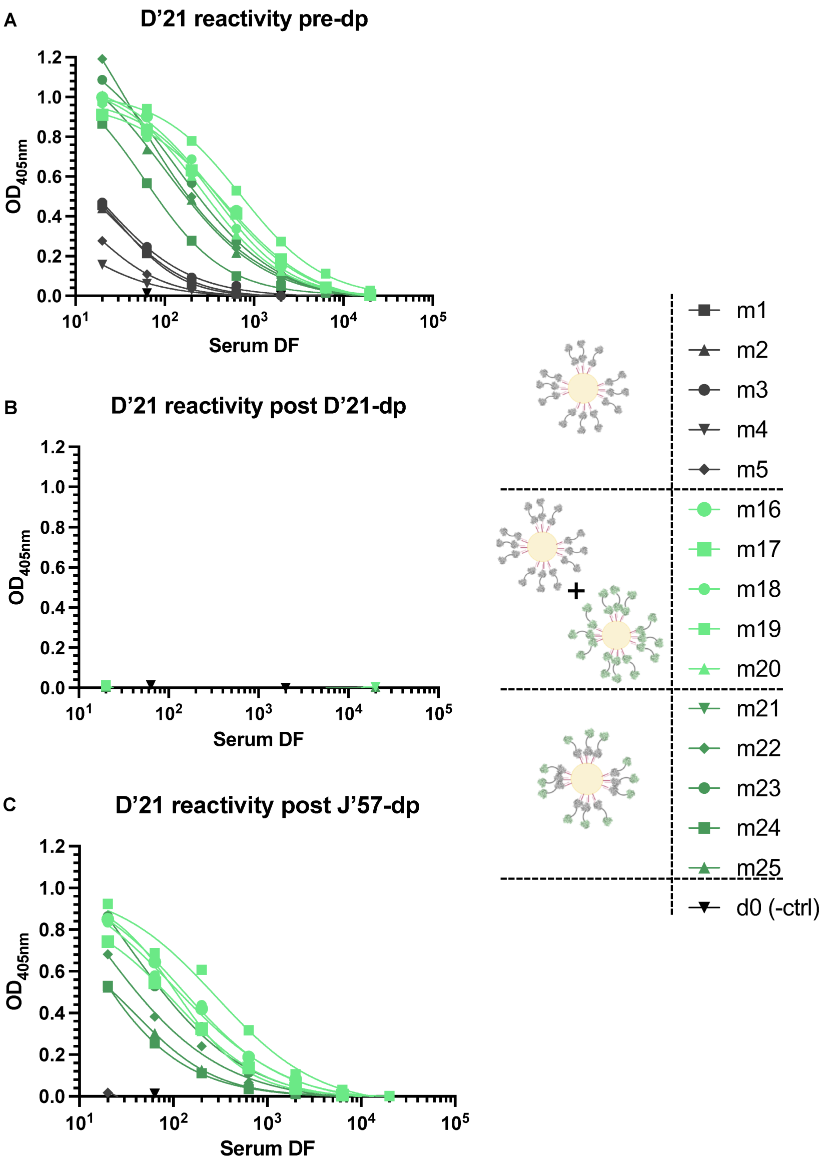


**fig. S11: Day 13 sera depletion studies for the D'21 NA. (A)** ELISA reactivity of the sera from mice m1-m5 and m16-m25 to D'21 NA prior to any strain depletions. Sera samples for mice m1-m5 and m16-m25 were depleted with D'21 NA antigen for any D'21 NA reactivity. D'21 NA-depleted sera samples were divided into two equal parts. One part was tested against D'21 NA, as shown in **(B)**, verifying that there was no remaining D'21 NA reactivity in the sera. (The other part of the D'21 NA-depleted sera was tested against J'57 NA.) **(C)** Sera from mice that were depleted for J'57 NA (m1-m5, m16-m25) were tested for remaining reactivity against D'21 NA.

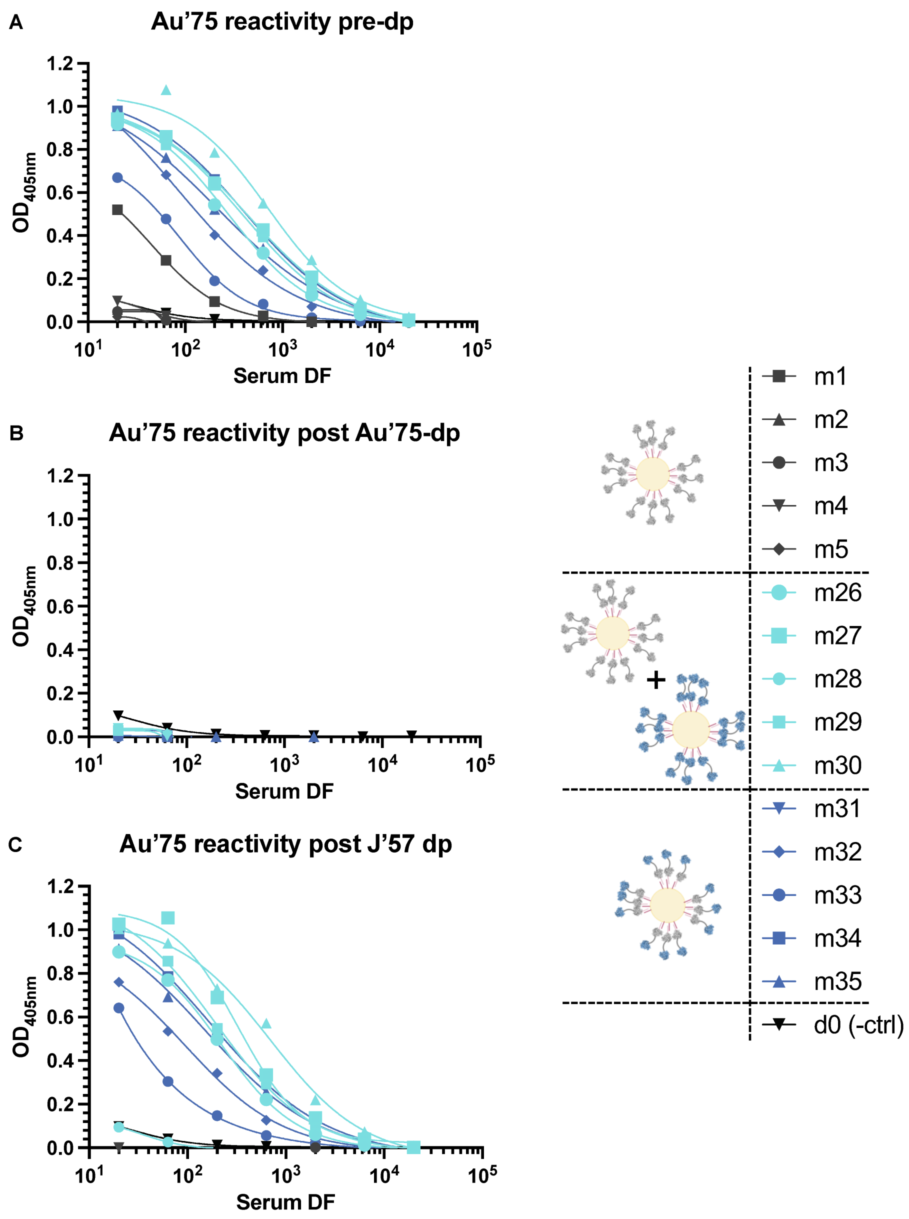


**fig. S12: Day 13 sera depletion studies for the Au’75 NA. (A)** ELISA reactivity of the sera from mice m1-m5 and m26-m35 to Au’75 NA prior to any strain depletions. Sera samples for mice m1-m5 and m26-m35 were depleted with Au’75 NA antigen for any Au’75 NA reactivity. One part was tested against Au’75 NA, as shown in **(B)**, verifying that there was no remaining Au’75 NA reactivity in the sera. (The other part of the Au’75 NA-depleted sera was tested against the J’57 NA strain). **(C)** Sera from mice that were depleted for J’57 NA (m1-m5, m26-m35) were tested for remaining reactivity against Au’75 NA.

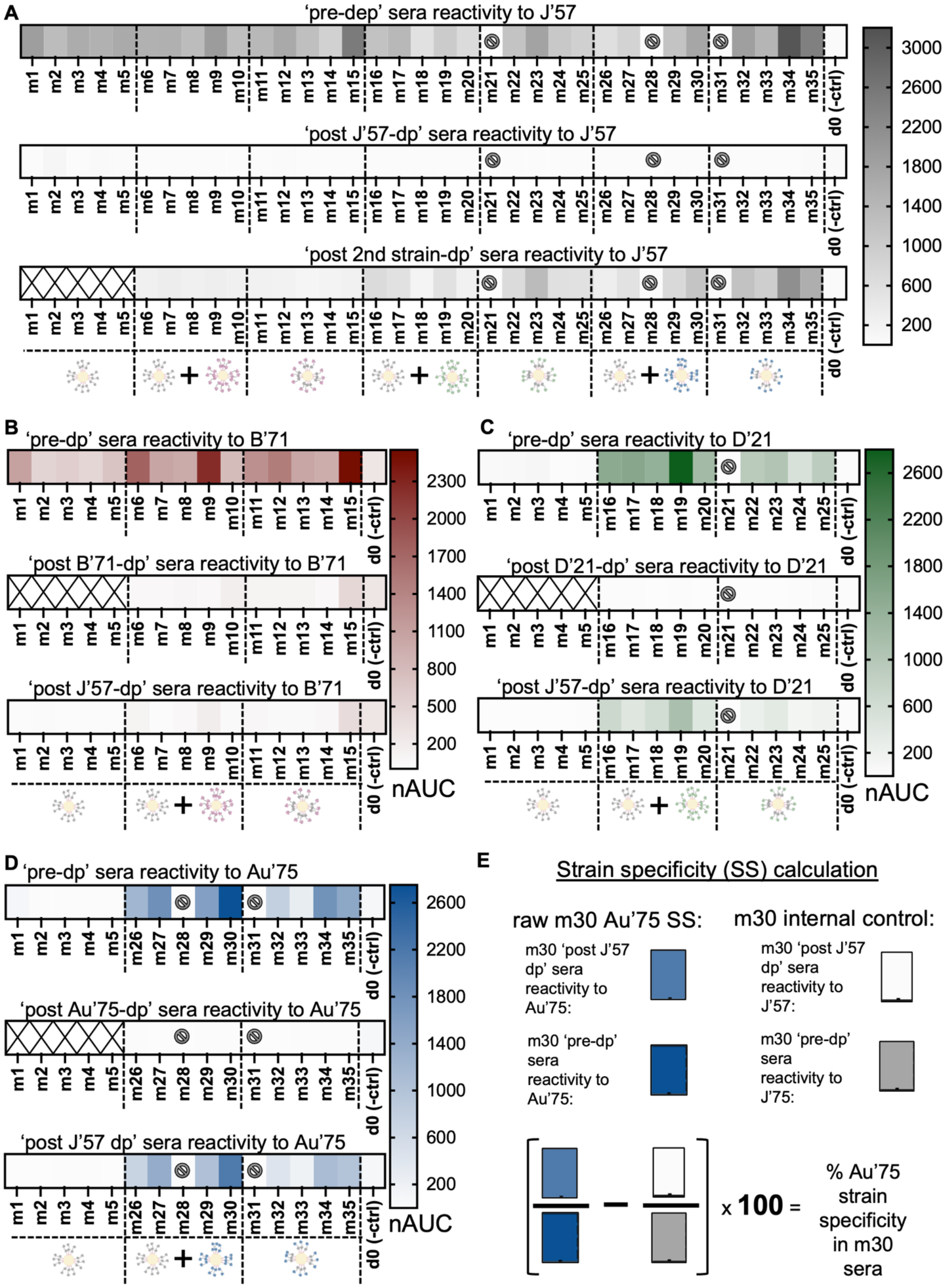

**fig. S13: Day 13 sera depletion normalized area under the curve (nAUC) values. (A)** nAUC heatmaps for the J'57 NA prior to any strain depletion (m1-m35), post J'57 NA depletion (m1-m35), and post B'71 or D'21 or Au'75 NA depletions (m6-m15, m16-m25, m26-m35, respectively). **(B)** nAUC heatmaps for the B'71 NA prior to any strain depletion (m1-m15), post B'71 NA depletion (m6-m15), and post J'57 NA depletion (m1-m15). **(C)** nAUC heatmaps for the D'21 NA prior to any strain depletions (m1-m5, m16-m25), post D'21 NA depletions (m16-m25), and post J'57 NA depletions (m1-m5, m16-m25). **(D)** nAUC heatmaps for the Au’75 NA prior to any strain depletions (m1-m5, m26-m35), post Au’75 NA depletion (m26-m35), and post J’57 NA depletion (m1-m5, m26-m35). **(E)** Example of how strain-specificity for a given sample was calculated. The serum from m30 was evaluated for reactivity to Au’75 NA prior to any NA depletions. After depleting the m30 serum for J’57 NA, the serum was tested for remaining reactivity to Au’75 NA. This represented the proportion of the serum that reacted solely to Au’75 NA. The remaining serum reactivity to Au’75 NA was divided by the starting serum reactivity to give a proportion of the serum that was strain-specific for Au’75 NA. To account for any residual cross-reactive serum antibodies in the eluted serum that were not removed, the remaining serum was also tested for reactivity to J’57 NA. That reactivity was divided by the starting serum reactivity to J’57 NA, and the proportion of remaining signal was subtracted from the Au’75 NA-specific proportion, thereby exacting stringent definitions for strain-specificity (*i.e.,* that proportion was used as the internal control for m30’s Au’75 NA strain-specificity). The difference in proportion was multiplied by 100 to obtain a percentage of the sera in m30 that was specific for strain Au’75 NA. Mice (m) m21, m28, and m31 did not respond to the prime immunization and were therefore removed from any statistical analyses.

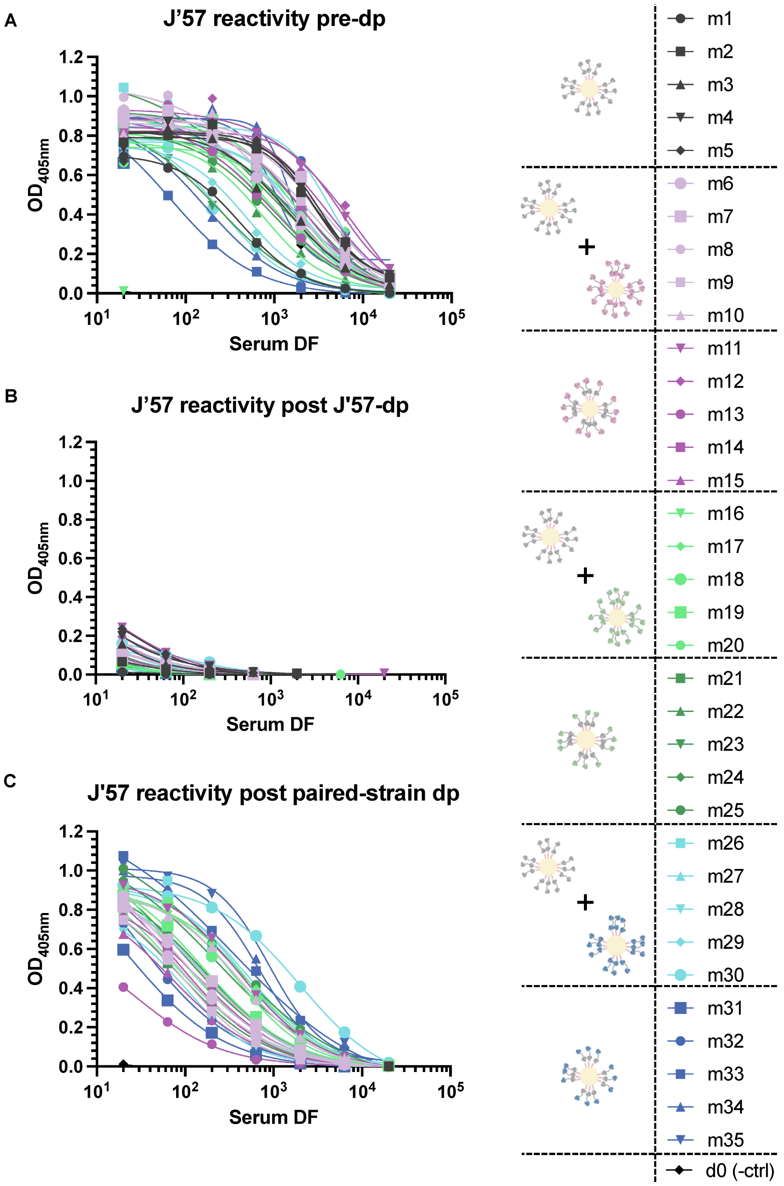


**fig. S14: Day 55 sera depletion data for J'57 NA.** (A) ELISA reactivity of the sera from all immunized mice to J'57 NA prior to any NA depletions. Sera samples from mice m1-m35 were depleted with J'57 NA. J'57 NA-depleted sera samples were divided into two equal parts. One part was tested against J'57 NA, as shown in **(B)**, verifying that there was no remaining J'57 NA reactivity in the sera. The other part of the J'57 NA-depleted sera was tested for reactivity against the remaining strains of NA. **(C)** Sera from mice that were depleted with B'71 NA (m6-m15), D'21 NA (m16-m25), and Au'75 NA (m26-m35) were tested for remaining reactivity against J'57 NA.

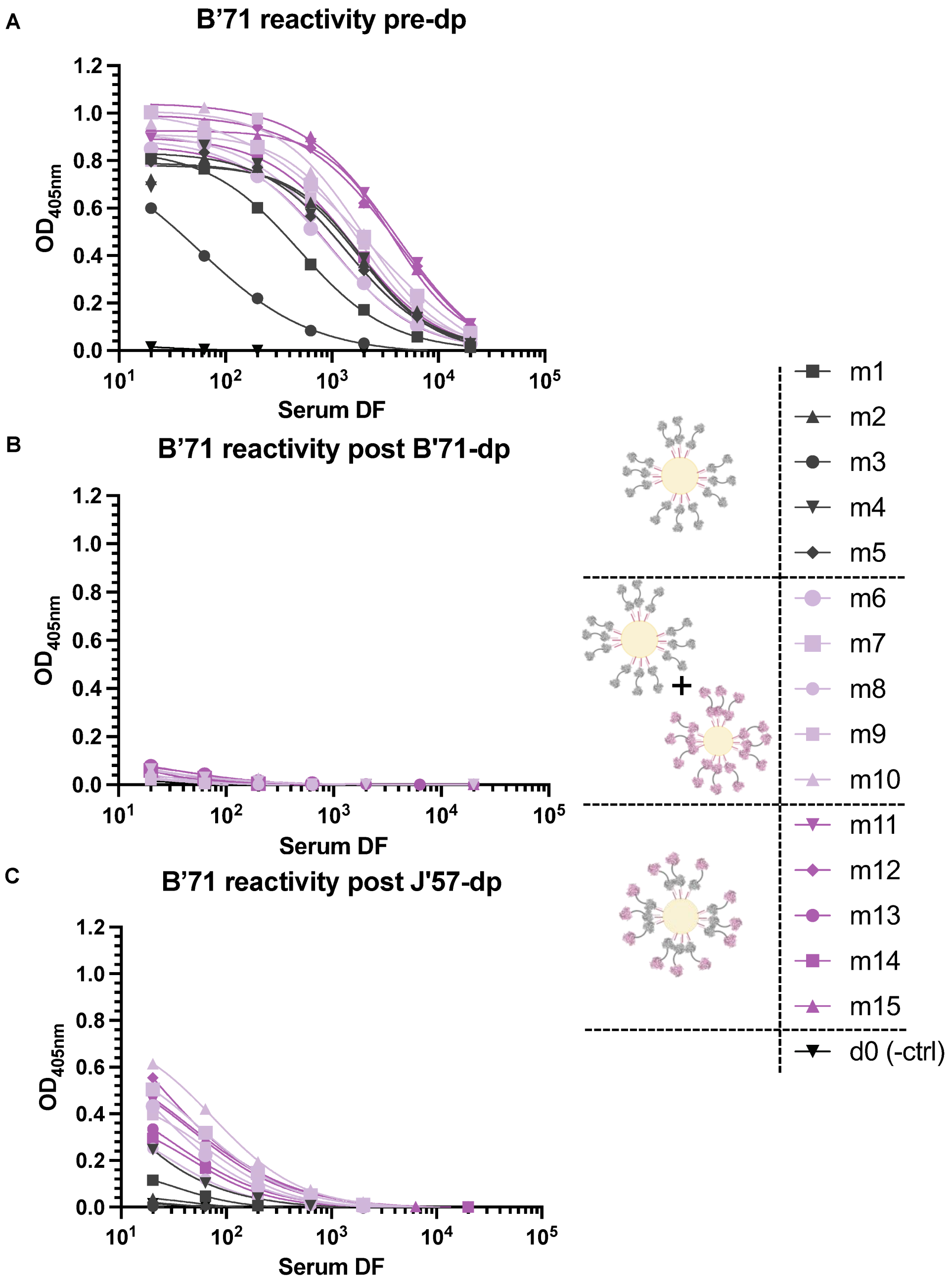


**fig. S15: Day 55 sera depletion data for B’71 NA. (A)** ELISA reactivity of the sera from mice m1-m15 to B’71 NA prior to any strain depletions. Sera samples for mice m1-m15 were depleted with B’71 NA antigen. B’71 NA-depleted sera samples were divided into two equal parts. One part was tested against B’71 NA, as shown in **(B)**, verifying that there was no remaining B’71 NA reactivity in the sera. (The other fraction of the B’71 NA-depleted sera were tested against J’57 NA.) **(C)** Sera from mice that were depleted for J’57 NA (m1-m15) were tested for remaining reactivity against B’71 NA.

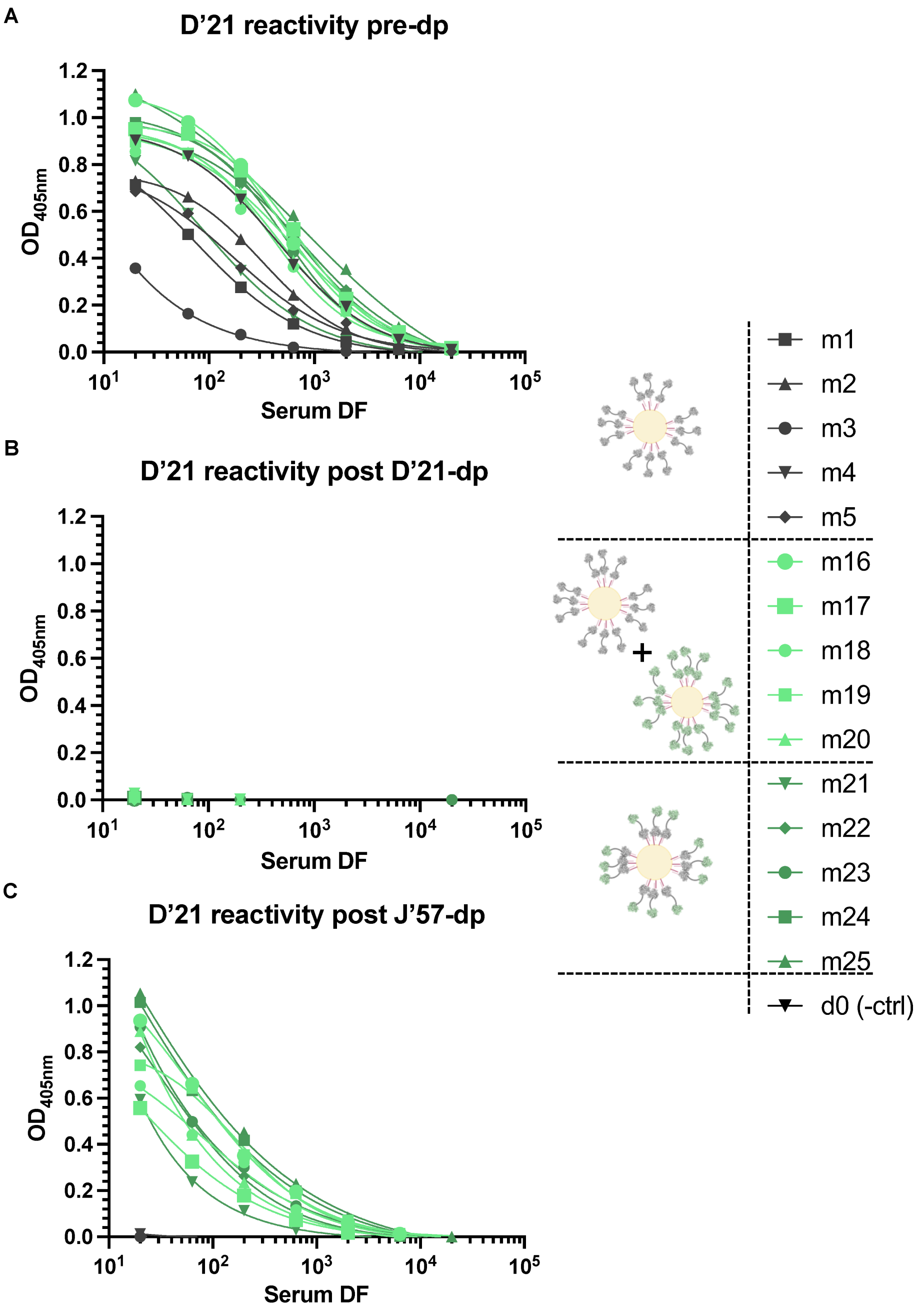


**fig. S16: Day 55 sera depletion data for D’21 NA. (A)** ELISA reactivity of the sera from mice m1-m5 and m16-m25 to D’21 prior to any NA depletions. Sera samples from mice m1-m5 and m16-m25 were depleted with D’21 NA. D’21 NA-depleted sera samples were divided into two equal parts. One part was tested against D’21 NA, as shown in **(B)**, verifying that there was no remaining D’21 NA reactivity. (The other fraction of the D’21-depleted sera were tested against J’57 NA.) **(C)** Sera from mice that were depleted with J’57 NA (m1-m5, m16-m25) were tested for remaining reactivity against D’21 NA.

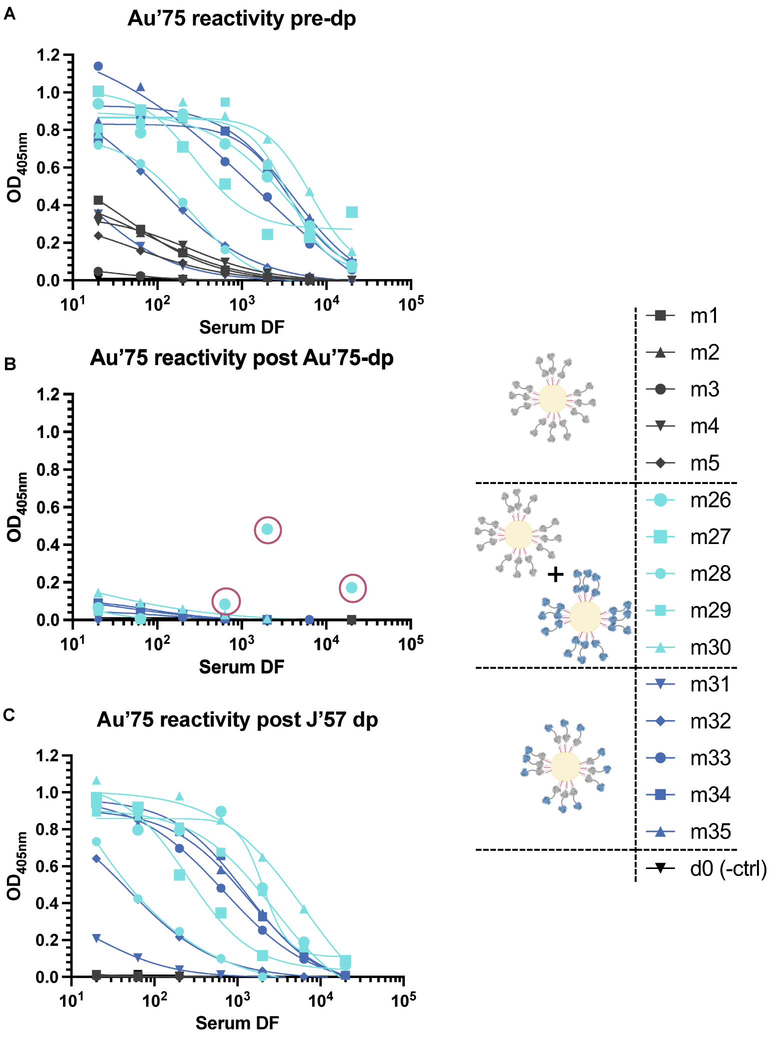


**fig. S17: Day 55 sera depletion data for Au’75 NA. (A)** ELISA reactivity of the sera from mice m1-m5 and m26-m35 to Au’75 prior to any NA depletions. Sera samples from mice m1-m5 and m26-m35 were depleted with Au’75 NA. Au’75 NA-depleted sera samples were divided into two equal parts. One fractopm was tested against Au’75 NA, as shown in **(B)**, verifying that there was no remaining Au’75 NA reactivity in the sera. The other part of the Au’75 NA-depleted sera was tested against J’57 NA. **(C)** Sera from mice that were depleted with J’57 NA (m1-m5, m26-m35) were tested for remaining reactivity against Au’75 NA. *Note: the three data points from m26 that circled red were removed from the nAUC analyses, as they contributed to a falsely elevated residual Au’75 reactivity that was clearly not biological.*

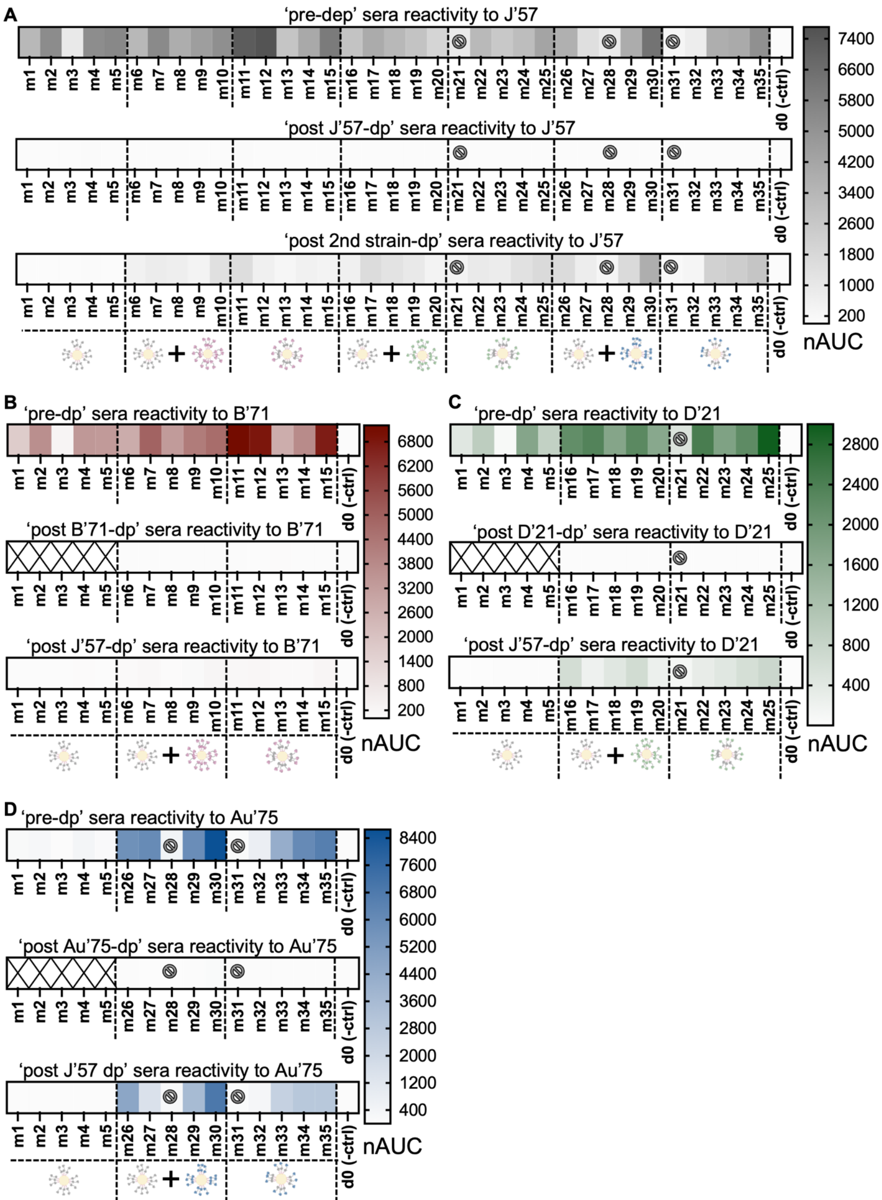


**fig. S18: Day 55 sera depletion normalized area under the curve (nAUC) values. (A)** nAUC heatmaps for J'57 NA prior to any strain depletion (m1-m35), post J'57 NA depletion (m1-m35), and post B'71 or D'21 or Au'75 NA depletions (m6-m15, m16-m25, m26-m35, respectively). **(B)** nAUC heatmaps for B'71 NA prior to any strain depletion (m1-m15), post B'71 NA depletion (m6-m15), and post J'57 NA depletion (m1-m15). **(C)** nAUC heatmaps for D'21 NA prior to any strain depletions (m1-m5, m16-m25), post D'21 NA depletions (m16-m25), and post J'57 NA depletions (m1-m5, m16-m25). **(D)** nAUC heatmaps for the Au'75 NA prior to any strain depletions (m1-m5, m26-m35), post Au'75 NA depletion (m26-m35), and post J'57 NA depletion (m1-m5, m26-m35). *Note: Mice (m) m21, m28, and m31 did not respond to the prime immunization and were therefore removed from any statistical analyses.*

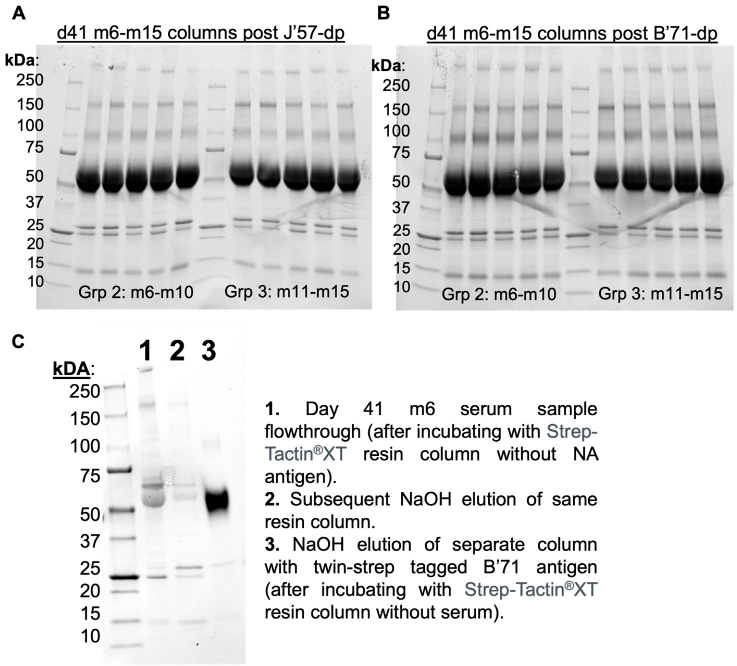


**fig. S19: Sample SDS PAGE analysis of NaOH-treated elution of d41 sera-depletion resin columns.** **(A)** Serum samples from m6-m15 were incubated with Strep-Tactin®XT resin that had been pre-incubated with excess StrepII-tagged J'57 NA. After the flowthrough sera was collected and washed with 100uL of PBS for use in depletion studies, the column was treated with NaOH and collected for SDS PAGE analysis (eluted samples were first concentrated using 10K MWCO). The band at 150kDa supports that J'57-binding antibodies were successfully depleted from the original serum sample. **(B)** serum samples from m6-m15 were incubated with Strep-Tactin®XT resin that had been pre-incubated with excess StrepII-tagged B'71 NA. After the flowthrough sera was collected and washed with 100uL of PBS for use in depletion studies, the column was treated with NaOH and collected for SDS gel analysis (samples were first concentrated using 10K MWCO). The band at 150kDa supports that B'71-binding antibodies were successfully depleted from the original serum sample. **(C)** Serum from m6 was incubated with Strep-Tactin®XT resin and flowthrough was collected after one hour and concentrated (column 1). The column was then immediately treated with NaOH (without a PBS wash), and the elution was concentrated for SDS PAGE analysis (column 2). Separately, StrepII-tagged B'71 was incubated for one hour and washed with PBS. The column was then treated with NaOH, and flowthrough was concentrated for SDS PAGE analysis (column 3). The nearly-absent band at 150kDa suggests that only residual antibodies from the sera remained in the column. For experimental data, residual antibodies were washed out of the column with 100uL of PBS.

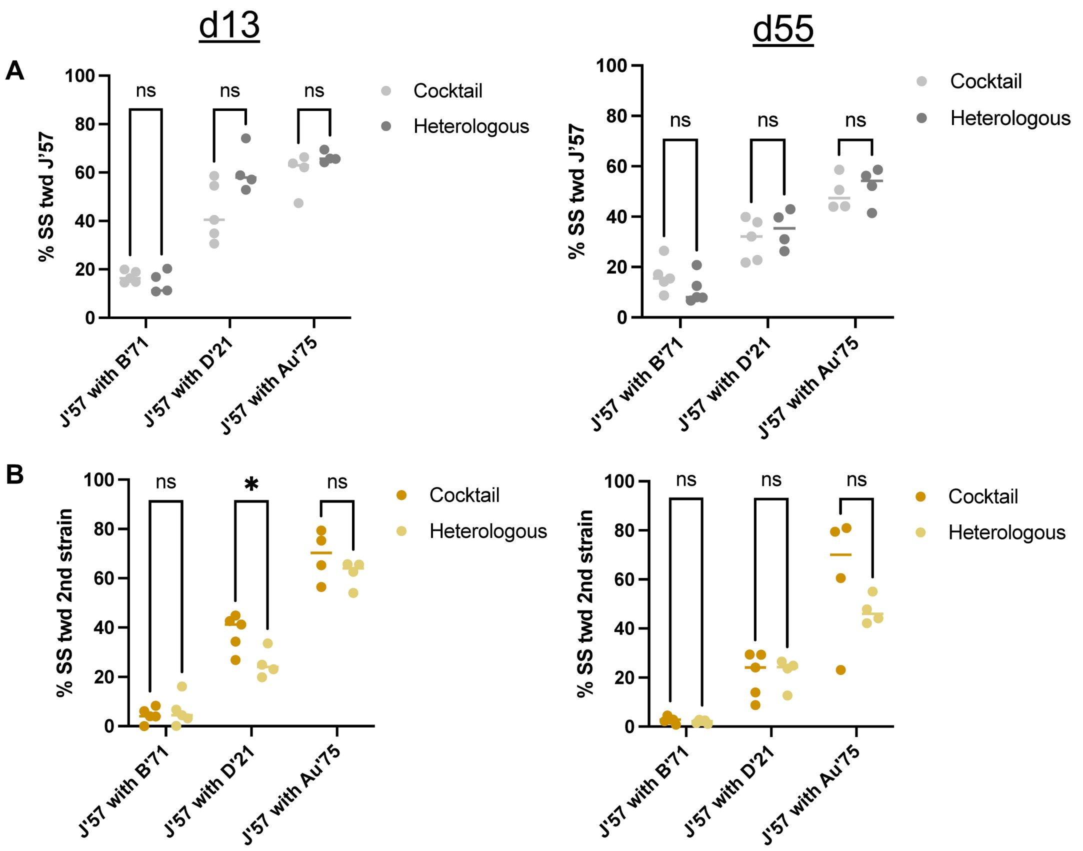


**fig. S20: Sera strain-specificity in the cocktail and heterologous NoaS NP groups. (A)** d13 and d55 comparison of J'57 NA strain-specificity within groups receiving homologous cocktails of J'57/J'57 NoaS NPs with either B'71/B'71 (group 2), D'21/D'21 (group 4), or Au'75/Au'75 (group 6), compared to the J'57 NA strain-specificity elicited in their counterpart heterologous NoaS NP groups of J'57/B71 (group 3), J'57/D'21 (group 5), and J'57/Au'75 (group 7). There were no statistically significant differences between groups 2 and 3, groups 4 and 5, and groups 6 and 7. Thus, data from groups 2 and 3 were pooled, groups 4 and 5 were pooled, and groups 6 and 7 were pooled. (B) Analogous strain specificity analyses for B'71, D'21, and Au'75 NAs. The only observed difference in strain-specificity for cocktail versus heterologous groups was J'57 with D'21 on d13 (p-value: *0.03). Thus, data from groups 2 and 3, groups 4 and 5, and groups 6 and 7 were evaluated collectively. Statistical significance was evaluated using Mann-Whitney tests.

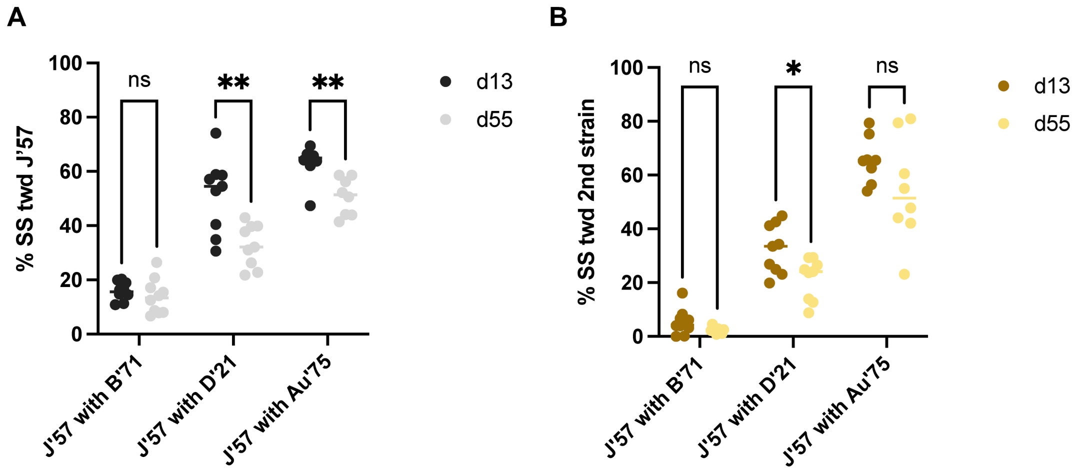


**fig. S21: Differences in strain-specificities before and after the boost immunization. (A)** Strain-specificity for J’57 NA was comparably low for the J’57/B’71 pairing on d13 and d55. However, strain specificity for J’57 NA decreased significantly on d55 compared to d13 for the groups containing J’57/D’21 (groups 4 and 5, p-value: **0.008) and J’57/Au’75 (groups 6 and 7, p-value: **0.003). **(B)** While the strain-specificity for B’71, D’21, and Au’75 NAs qualitatively decreased on d55 compared to d13, the decrease was only statistically significant for strain-specificity to D’21 (*i.e.,* groups 4 and 5, p-value: *0.04). Statistical significance was evaluated using Mann-Whitney tests.

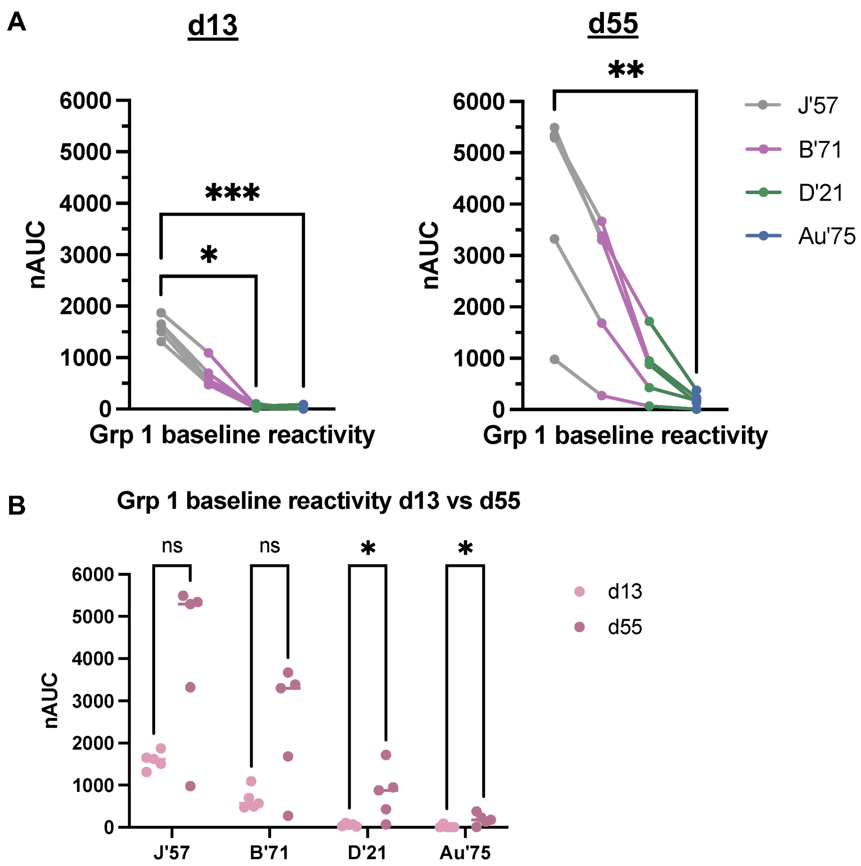


**fig. S22: Development of NA breadth in group 1. (A)** Baseline group reactivity of the group 1 (grp1) J’57/J’57 control to each strain. Group 1 had significantly lower reactivity to D’21 NA (p-value: *0.0234) and Au’75 NA (p-value: ***0.0009) compared to the J’57 NA on d13. By d55, group 1 reactivity increased to each strain, and only reactivity to Au’75 was significantly lower than the J’57 strain (p-value: **0.0046). Statistical significance was assessed using Kruskal-Wallis with Dunn correction for multiple comparisons. **(B)** Group 1 baseline reactivity for each NA strain was assessed at d13 and d55. Statistical significance was noted for D’21 NA (p-value: *0.02) and Au’75 NA (p-value: *0.02), which increased significantly on d55 compared to d13. Statistical significance was assessed using Mann-Whitney tests.

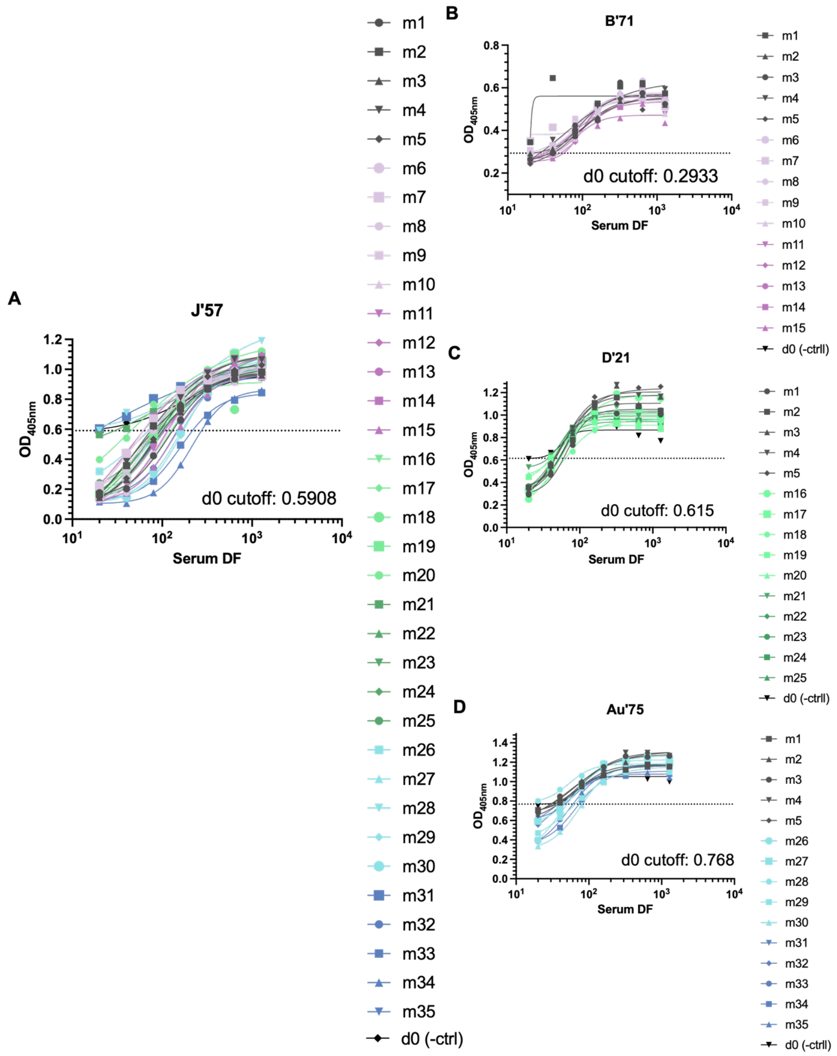


**fig. S23: Day 20 maximum dilution factor of inhibition data.** **(A)** Serial dilutions of sera from m1-m35 were premixed with 2x the $EC_{50}$ concentration value of J'57 NA tetramers and tested for NA enzymatic activity in the ELLA assay. The maximum dilution factor of inhibition (MDFI) was defined per mouse sera as the maximum dilution factor of a mouse sera that inhibited the activity of the J'57 NA tetramer better than the highest concentration of the d0 negative control serum. (The $OD_{405nm}$ at the highest dilution of d0 sera was used as the cutoff for the MDFIs for the remaining sera samples.) **(B)** Serial dilutions of sera from m1-m15 were premixed with 2x the $EC_{50}$ concentration value of B'71 NA monomers and evaluated for NA enzymatic activity inhibition in the ELLA assay. The MDFI was defined per mouse as the maximum dilution factor of a mouse sera that inhibited the activity of the B'71 NA better than highest concentration of the d0 negative control serum. **(C)** Serial dilutions of sera from m1-m5 and m16-m25 were premixed with 2x the $EC_{50}$ concentration value of D'21 NA tetramers and tested for NA enzymatic activity in the ELLA assay. The MDFI was defined per mouse as the maximum dilution factor of sera that inhibited the activity of the D’21 NA tetramer better than highest concentration of the d0 negative control serum. **(D)** Serial dilutions of sera from m1-m5 and m26-m35 were premixed with 2x the $EC_{50}$ concentration value of Au’75 NA tetramers and tested for NA enzymatic activity in the ELLA assay. The MDFI was defined per mouse as the maximum dilution factor of mouse sera that inhibited the activity of the Au’75 NA tetramer better than the highest concentration of the d0 negative control serum.

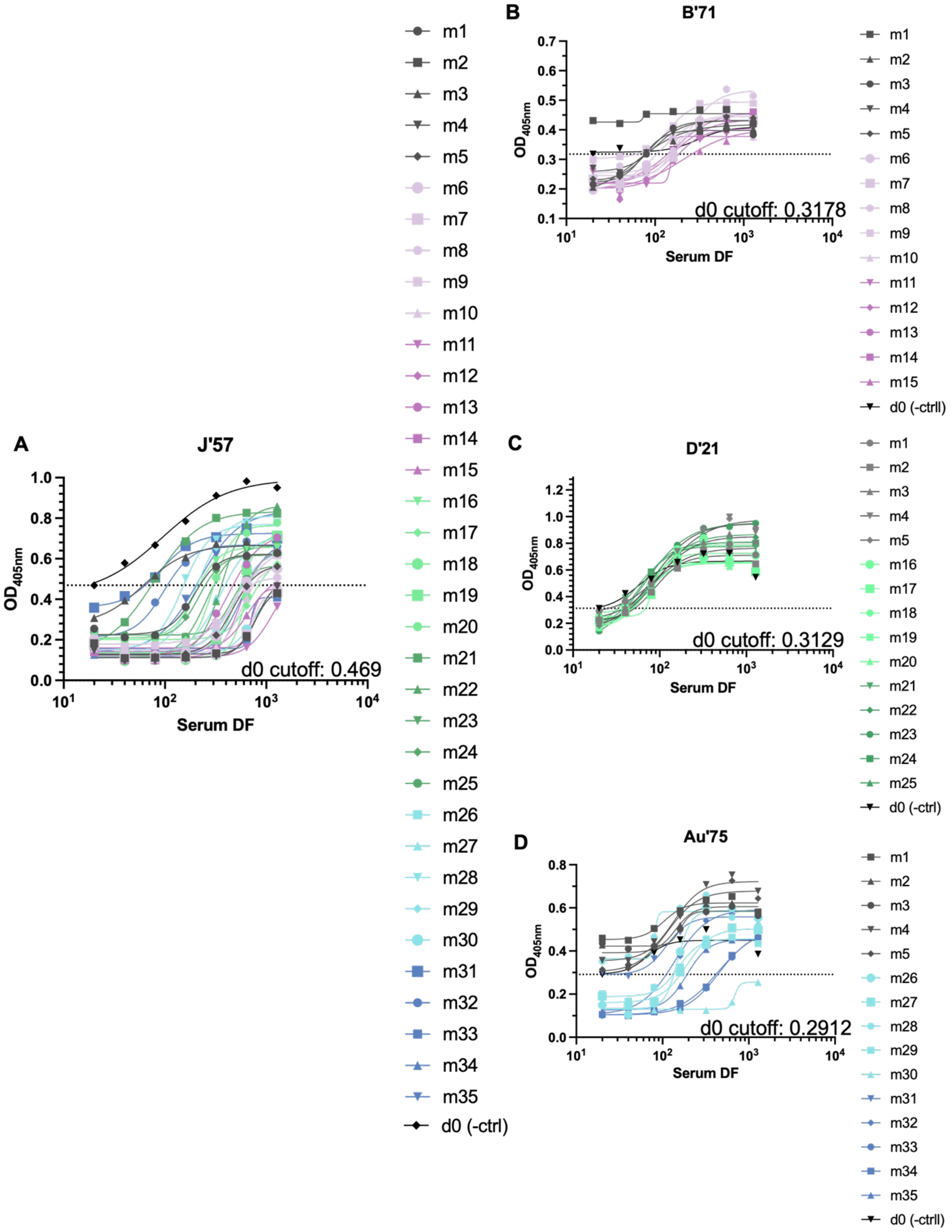


**fig. S24: Day 62 maximum dilution factor of inhibition data. (A)** Serial dilutions of sera from m1-m35 were premixed with 2x the $EC_{50}$ concentration value of J'57 NA tetramers and tested for NA enzymatic activity in the ELLA assay. The maximum dilution factor of inhibition (MDFI) was defined per mouse sera as the maximum dilution factor of a mouse sera that inhibited the activity of the J'57 NA tetramer better than the highest concentration of the d0 negative control serum. (The $OD_{405nm}$ at the highest dilution of d0 sera was used as the cutoff for the MDFIs for the remaining sera samples.) **(B)** Serial dilutions of sera from m1-m15 were premixed with 2x the $EC_{50}$ concentration value of B'71 NA tetramers and evaluated for NA enzymatic activity inhibition in the ELLA assay. The MDFI was defined per mouse as the maximum dilution factor of a mouse sera that inhibited the activity of the B’71 NA better than highest concentration of the d0 negative control serum. **(C)** Serial dilutions of sera from m1-m5 and m16-m25 were premixed with 2x the $EC_{50}$ concentration value of D’21 NA tetramers and tested for NA enzymatic activity in the ELLA assay. The MDFI was defined per mouse as the maximum dilution factor of sera that inhibited the activity of the D’21 NA tetramer better than highest concentration of the d0 negative control serum. **(D)** Serial dilutions of sera from m1-m5 and m26-m35 were premixed with 2x the $EC_{50}$ concentration value of Au’75 NA tetramers and tested for NA enzymatic activity in the ELLA assay. The MDFI was defined per mouse as the maximum dilution factor of mouse sera that inhibited the activity of the Au’75 NA tetramer better than the highest concentration of the d0 negative control serum.

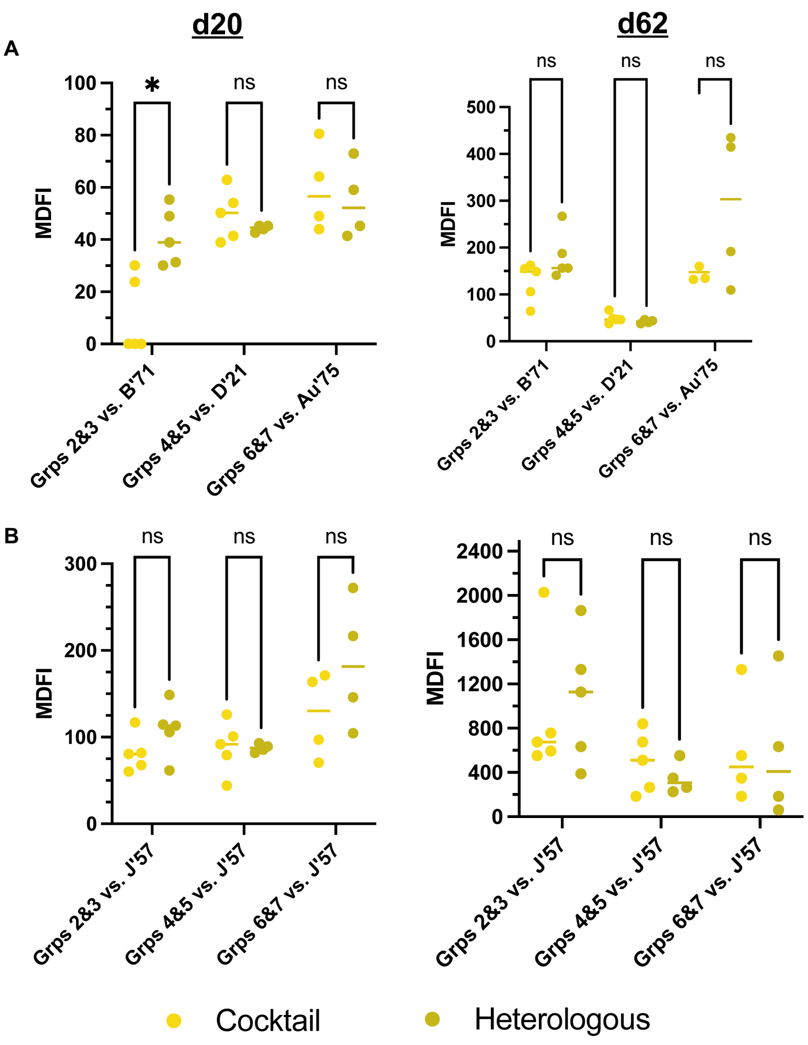


**fig. S25: Contrast of MDFIs of homologous cocktail group vs. their heterologous NP group. (A)** Comparison of the MDFIs of group 2 versus group 3 with B'71 NA activity inhibition, group 4 versus group 5 with D'21 NA activity inhibition, and group 6 versus group 7 to Au'75 NA activity inhibition on d20 and d62, respectively. Except for group 2 versus group 3 on d20 (p-value: *0.03), the cocktail vs. heterologous groups were statistically comparable, and the data were therefore pooled to reflect differences based on immunized strains rather than NP presentations. **(B)** Comparison of the MDFIs of the cocktail vs. their respective heterologous groups in their ability to inhibit J'57 NA tetramer activity. All cocktail vs. heterologous groups were comparable, and the data was therefore pooled to reflect differences based on immunized strains rather than NP presentations. *Note: m30 in group 6 had an outlier value of 1280 on d62, so it is not shown in the graph in **(A)** despite being included in the analyses.*

**REFERENCES**


1. Cho, K. J. *et al.* The Crystal Structure of Ferritin from *Helicobacter pylori* Reveals Unusual Conformational Changes for Iron Uptake. *J. Mol. Biol.* **390**, 83–98 (2009).
2. Zakeri, B. *et al.* Peptide tag forming a rapid covalent bond to a protein, through engineering a bacterial adhesin. doi:10.1073/pnas.1115485109/-/DCSupplemental.
3. Klein, J. S. *et al.* Design and characterization of structured protein linkers with differing flexibilities. in vol. 27 325–330 (Oxford University Press, 2014).
4. Stadlbauer, D. *et al.* Broadly protective human antibodies that target the active site of influenza virus neuraminidase.
5. Lederhofer, J. *et al.* Protective human monoclonal antibodies target conserved sites of vulnerability on the underside of influenza virus neuraminidase. *Immunity* **57**, 574-586.e7 (2024).
6. Zhu, X. *et al.* Structural Basis of Protection against H7N9 Influenza Virus by Human Anti-N9 Neuraminidase Antibodies. *Cell Host Microbe* **26**, 729-738.e4 (2019).
7. Lei, R. *et al.* Leveraging vaccination-induced protective antibodies to define conserved epitopes on influenza N2 neuraminidase. *Immunity* **56**, 2621-2634.e6 (2023).
